# Rapid microbiology instrumentation for high-accuracy quantitative water safety assessments in remote and low-resource locations

**DOI:** 10.1101/2025.06.13.659038

**Authors:** Dan E. Angelescu, David Abi Saab, Emilie Vilana, Mathias Haguet, Franziska Knoche, Nicolas Caradot, Wolfgang Seis

## Abstract

Microbial contamination of surface waters remains a significant public health concern, particularly in low- and middle-income countries (LMICs) and urban settings affected by combined sewer overflows (CSOs). Standard culture-based methods for monitoring fecal indicator bacteria (FIB)— including membrane filtration (MF) and most probable number (MPN) assays—typically require laboratory infrastructure, do not provide same-day results, and systematically underestimate total FIB loads due to their inability to accurately quantify aggregate-bound bacteria.

This study evaluates ALERT, an automated rapid method for culturable FIB quantification, as a viable field-deployable alternative to conventional laboratory techniques. ALERT employs a whole-sample approach that comprehensively captures both planktonic and aggregate-bound *E. coli*, thereby addressing critical limitations of traditional methods. Three instrument variants were assessed: the in-situ ALERT System V2, the portable ALERT LAB, and the handheld ALERT One, designed for single-sample analysis in resource-limited environments.

Comparative field and laboratory trials conducted in Germany, France, and the UK demonstrate that ALERT delivers accuracy and precision equivalent to MPN methods, while providing faster results directly in the field and enabling high-frequency sampling. Use cases included storm event monitoring, CSO impact evaluation, and on-site validation and optimization of point-of-use water potabilization treatments. ALERT One was also successfully used by citizen scientist volunteers, highlighting its suitability for decentralized community-based monitoring.

By enabling rapid, accurate, and comprehensive microbial water quality assessments, ALERT qualifies as a robust field alternative to laboratory testing. It supports enhanced water safety, empowers community-based surveillance and public engagement, and facilitates data-driven decision-making in both LMICs and high-income countries.

## 1 Introduction

The presence of fecal pathogens in water poses a significant global health risk, with contaminated drinking water, food chain pollution, and recreational exposure driving waterborne disease transmission. According to the World Health Organization (WHO, 2022a), microbiological contamination is responsible for over 1.5 million diarrheal-related deaths annually, making it one of the leading causes of mortality among young children worldwide (UNICEF, 2023). Recent estimates indicate that approximately 4.4 billion people in low- and middle-income countries (LMICs) lack access to safely managed drinking water (Greenwood, 2024), primarily due to fecal contamination as evidenced by the frequent detection of *Escherichia coli* (*E. coli*) bacteria. This risk is particularly severe in Southern Asia and sub-Saharan Africa, where reliance on unsafe water sources remains widespread. Given that fecal contamination is a primary driver of epidemics and diarrheal diseases, systematic microbiological water safety surveillance is crucial for public health protection— especially in LMICs where centralized treatment and distribution infrastructures are often absent. Yet, the resource-intensive nature of traditional monitoring methods has led to a critical data gap, with microbiological water quality data missing for large percentages of the population outside Europe and North America (UNICEF-WHO, 2023).

In developed nations, significant infrastructure advancements and stringent water safety measures— including quantitative microbial risk assessments (QMRA)—have resulted in notable reductions in drinking water-related disease outbreaks (WHO, 2022b). However, waterborne diseases still impose a considerable public health and economic burden. For example, in North America, fecal pathogens from contaminated irrigation sources and recreational waters contribute to annual costs amounting to billions of dollars (Marshall et al., 2020; Spalding et al., 2023; DeFlorio-Barker et al., 2018). Furthermore, the ongoing climate crisis, characterized by increasing weather extremes and global precipitation variability (Zhang, 2024), is exacerbating challenges related to water availability and quality (IPCC, 2021). This situation has led to a growing dependence on alternative water sources, such as recycled wastewater, which introduces additional health risks (Drechsel et al., 2022).

Epidemiological studies have established links between the presence and concentration of viable, culturable fecal indicator bacteria (FIB)—such as *E. coli* and enterococci—and the incidence of waterborne disease in recreational waters (Cabelli, 1983; Dufour, 1984; Kay et al., 2004). As a result, quantitative *E. coli* measurements have become central for assessing the microbiological quality of surface waters, treated municipal wastewater, but also of drinking water (WHO, 2022b). In most jurisdictions, regulations require that *E. coli* (or, in some cases, thermotolerant coliforms) be non-detectable in 100 mL of drinking water. While developed countries with well-maintained, disinfected distribution networks often rely on presence/absence tests for routine monitoring (quantitative testing being required for source investigations and raw water monitoring), many LMICs face widespread, persistent fecal contamination; for instance, in some African nations, over 80% of the population relies on unsafe water sources (WHO, 2021). In such contexts, presence/absence tests are inadequate, making quantitative *E. coli* measurements critical for prioritizing safer water sources to reduce waterborne disease-related mortality (UNICEF, 2019). To support risk-based water safety assessments, WHO has established a risk level classification system based on *E. coli* concentrations (WHO, 2022b).

The large-scale deployment of accurate microbiological water quality testing is currently hindered by several factors. Traditional culture-based methods—such as membrane filtration (MF) with agar plating and most-probable-number (MPN) assays—require representative sampling, rapid refrigerated transport, serial dilutions, and, at times, confirmatory tests. Each step introduces potential sources of human error and sample degradation but also important delays, resulting in a time-to-result (TTR) of at least 24 hours; in practice, results are often available only several days after sample collection. In many LMIC locations and in remote areas worldwide, maintaining sample integrity can be challenging and laboratory facilities may be unavailable, rendering microbiological testing impossible.

A further limitation to accurate monitoring comes from certain intrinsic limitations of current laboratory methods. Conventional culture-based techniques assume fully homogenized samples—a condition that is frequently unmet in the presence of bacteria attached to particles. Under such circumstances, these methods cannot differentiate between free-floating individual bacteria (planktonic count) and those aggregated in larger numbers to sediment or fecal particles (USEPA, 2012b; Jarvis et al., 2010), leading to major potential underestimation of the total (comprehensive) FIB load and consequent risk misclassification. The substantial presence of aggregate-bound bacteria and the resulting discrepancies were demonstrated in different contexts, including for the Seine river at the location of the 2024 Olympic aquatic events (Angelescu *et al*., 2024a).

Cost-effectiveness further complicates routine water quality testing. In many LMIC settings safe water distribution networks are unavailable, while financial constraints limit access to routine testing. In such settings, decentralized water treatment solutions—such as iJal water ATMs in rural India (Sewak, 2020) and various household treatment methods (WHO, 2025)—are on the rise. However, monitoring the real-world effectiveness of these solutions under variable field conditions remains both challenging and prohibitively expensive. Even in developed nations, large-scale FIB monitoring programs incur substantial costs, particularly in remote regions. In response, community-based monitoring initiatives have emerged as a promising approach to collect water quality data that would otherwise be too costly or logistically challenging to obtain, including in LMICs (UNICEF, 2019). Unfortunately, until today community-based “citizen science” projects have predominantly focused on physical and chemical parameters, with very limited inclusion of FIB monitoring despite the associated public health importance (Ramirez et al., 2023).

To overcome these challenges, there is a pressing need for new accurate *E. coli* quantification technologies that are portable, field-deployable, and independent of specialized laboratory facilities or highly trained personnel. For successful implementation in LMICs, these methods must be easy to use, enable simple result reporting, and deliver rapid, same-day outcomes—ideally within a few hours—to enable prompt response and mitigation. Moreover, new testing methods must overcome the limitations of traditional culture-based techniques by comprehensively capturing both free-floating and aggregate-bound *E. coli*. Developing cost-effective, automated on-site *E. coli* testing instrumentation has the potential to revolutionize water quality monitoring and significantly expand the reach of microbiological testing in LMICs and remote areas worldwide, ultimately saving lives from preventable waterborne diseases.

This study assesses the potential of Fluidion^®^ ALERT rapid *E. coli* quantification technology as a viable alternative to conventional laboratory-based monitoring methods for cost-effective, on-site microbiological risk assessments in LMICs and remote locations. We conduct a thorough metrological evaluation of the technology and demonstrate its utility across three key applications: i. Monitoring contaminated surface waters in urban and peri-urban settings; ii. Enabling community-based monitoring through citizen science initiatives; and iii. Testing drinking water supplies and point-of-use treatment systems developed for LMIC environments and crisis response situations.

We start by presenting a direct metrological evaluation of portable and *in-situ* ALERT instruments, using laboratory-spiked samples and avoiding the challenges posed by presence of FIB aggregates. This part of our study allowed evaluating precision, linearity, and offset relative to standard MPN measurements in controlled conditions. We also discuss a straightforward procedure for adapting the master ALERT calibration to account for local water-specific matrix effects, thereby eliminating potential site-specific measurement bias.

We then present results from several field deployment case studies that were conducted to assess the performance of three distinct ALERT instrument versions (*in-situ*, portable and handheld) under real-world conditions in urban and peri-urban settings. Several months of time-series data of ALERT *E. coli* concentrations were collected from locations in the Seine River in Paris, France, and the Spree Canal in Berlin, Germany, using portable and *in-situ* ALERT instruments concurrently with side-by-side MPN laboratory measurements. The deployment of handheld ALERT instruments is exemplified through a community-monitoring project of the Thames River around Henley-on-Thames, U.K. We discuss how these field studies demonstrate the ability of ALERT technology to capture both short- and long-term dynamics of microbiological pollution during short-term pollution (STP) events caused by storms or combined sewer overflows (CSOs), and evaluate the implications of aggregate-bound fecal indicator bacteria for surface water analysis. We further discuss the feasibility of substituting laboratory-based analyses with ALERT technology for recreational water monitoring, which is demonstrated by verifying agreement with current U.S. and European approved regulatory approaches.

Finally, we demonstrate how ALERT technology can address two critical applications for LMICs: dosage optimization for point-of-use potabilization treatments, and on-site assessment of drinking water microbiological quality. Specifically, we test different concentrations of a widespread on-site flocculation-coagulation-chlorination water treament on microbiologically-contaminated urban surface waters, and assess their respective efficacy on free-floating (planktonic) *E. coli versus* aggregate-bound forms. We then present results from side-by-side independent laboratory testing, performed on behalf of the WHO, and discuss the ability of ALERT technology to rapidly and accurately quantify fecal contamination—and the corresponding risk class—in potentially unsafe drinking water sources in LMICs.

## 2 Materials and Methods

Several distinct studies were undertaken to evaluate the capabilities of ALERT instrumentation to provide rapid and accurate measurements of *E. coli* concentration directly in the field, including in remote or LMIC contexts. These included laboratory-based metrology testing using surface water samples spiked with wastewater treatment plant (WWTP) effluent, side-by-side time-series data collection from multiple field deployments in different canal and river locations (including through community-based citizen science), and field evaluation and optimization for a widespread point-of-use water treatment solution. Specific focus, including new microscopic evidence, was placed on the presence of *E. coli* aggregated onto fecal particles and their implications in terms of recreational water quality classification and point-of-use water treatment efficacy. The WHO performed an independent verification of ALERT technology for drinking water analysis as part of the UNICEF Rapid Water Quality Testing Initiative (UNICEF, 2025), with some of the relevant methodology aspects summarized below and detailed presentation of the respective protocols available in the published verification report (WHO-UNICEF, 2022).

### 2.1 Microbiology procedures

#### 2.1.1 Laboratory MPN *E. coli* measurements

All MPN measurements for the Berlin and the Ablon-sur-Seine studies were performed using miniaturized 96-well plate (ISO, 1998) at approved laboratories, using the bathing water (Berlin) or surface water (Ablon-sur-Seine) dilution protocols; confidence intervals were not reported.

The IDEXX Colilert Quantitray-2000 system was used to perform all MPN measurements for the WHO verification protocol (WHO-UNICEF, 2022), following standardized protocols (ISO, 2012).

#### 2.1.2 ALERT *E. coli* measurements

##### 2.1.2.1 ALERT methodology

The ALERT FIB quantification method used in this study implements a whole-sample culture-based approach with real-time enzymatic detection (Angelescu et al. 2019). The sample is incubated in the presence of a bioreagent containing a specific growth medium. This feature is central to the ALERT measurement process and restricts the measurement to viable and culturable bacteria, such specificity being required by water quality regulations worldwide. The further contains the enzymatic substrate 4-methylumbelliferyl-β-D-glucuronide (MUG), which is progressively metabolized by *E. coli* into the fluorescent by-product methylumbelliferone (MUF). Real-time monitoring of the MUF fluorescence in the whole sample during the bacterial culture process allows measuring all *E. coli* present in the sample, regardless of their free or particle-attached status, through a quantification mechanism that is similar to the one used in quantitative polymerase chain reaction (qPCR) methods (Angelescu *et al*., 2024a). Raw samples were analyzed to obtain the comprehensive (or total) *E. coli* count, including bacteria that are attached individually or in larger numbers to larger fecal or sediment particles, whereas 5 µm-filtered samples were analyzed to obtain the planktonic (or free-floating) *E. coli* count (on days with high turbidity, multiple filters were required due to progressive filter clogging). The 5 µm threshold was selected to allow only free bacteria and very small aggregates to pass through, while retaining larger aggregates.

##### 2.1.2.2 ALERT instrumentation used

The ALERT quantification method is deployed across three complementary device formats: the ALERT System V2 (a fully automated, cloud-connected, in-situ analyzer), the ALERT Lab (a portable, cloud-connected six-sample analyzer), and ALERT One (a cost-effective, handheld single-sample device).

The ALERT System V2 performs up to seven autonomous measurements using preloaded sterile disposable cartridges containing liquid bioreagent. Sampling, preparation, and analysis are fully automated and can be scheduled or triggered remotely. Once all cartridges are used, the system requires on-site cartridge replacement as part of routine maintenance.

For ALERT Lab and ALERT One, the user manually mixes a measured volume of water (25 or 50 mL) with liquid bioreagent before initiating the measurement. For low-resource drinking water testing, the bioreagent can also be supplied in powder form within a disposable vial, requiring simple dissolution by stirring. ALERT Lab is remotely controlled and enables parallel testing in six independent reaction chambers, while ALERT One provides a single reaction chamber with integrated local control via screen and buttons, performing all quantification steps internally.

Measurement results and automatically-generated PDF reports—including quality control elements such as raw optical signal curves, incubation temperature logs, and water temperature data—were accessed via the Fluidion Data Analytics cloud platform for ALERT Lab and ALERT System V2, or via a computer application for ALERT One. Figure 1 depicts the three devices in field use, while Table 1 summarizes the ALERT devices employed in the different components of this study.

**Figure 1.**
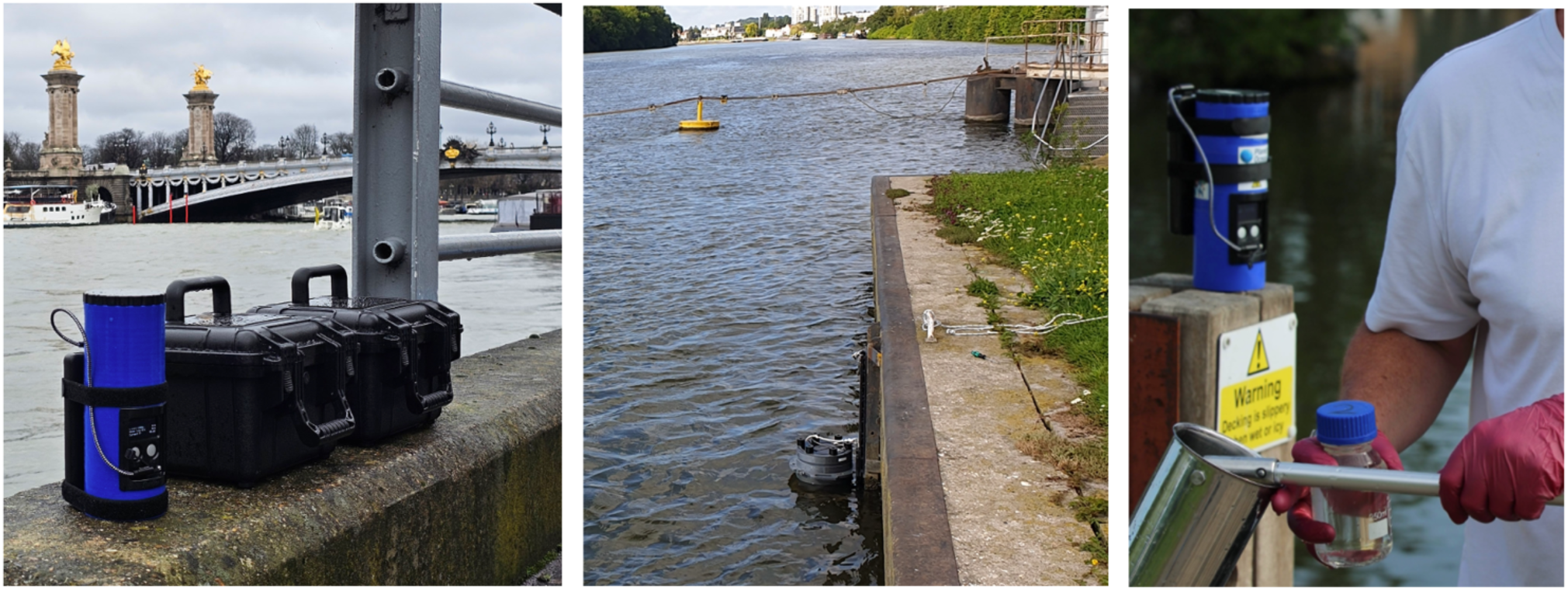
Left: ALERT One and ALERT Lab instruments on the banks of the Seine river in Paris, at the 2024 Olympic venue. Middle: ALERT System V2 deployed at Ablon-sur-Seine. Right: Citizen scientist sampling and using the ALERT One instrument at Henley-on-Thames (photo courtesy of the HoT Water group).

**Table 1.** Study names, locations and instrumentation types used in this study.

| Study type (water matrix) | Location | Instruments used |
| --- | --- | --- |
| **Metrology study (surface, canal)** | Berlin (Germany) | ALERT Lab<br>ALERT System V2 |
| **Side-by-side time series (surface, canal)** | Spree Canal Berlin (Germany) | ALERT Lab |
| **In-situ side-by-side time series (surface, river)** | Seine River, Ablon-sur-Seine (France) | ALERT System V2 |
| **Citizen-science time series (surface, river)** | Thames River, Henley (U.K.) | ALERT One |
| **Point-of-use water treatment solution evaluation (surface, river)** | Alfortville (France) | ALERT Lab<br>ALERT One |
| **Drinking water quality evaluation** | WHO KWR laboratory, Netherlands | ALERT Lab |

### 2.2 Sample preparation for metrology study

Metrology studies were conducted using the ALERT Lab and ALERT System V2 to quantify the offset and precision of these devices compared to the laboratory reference method ISO 9308-3 under controlled conditions. To this end repeated measurements of the ALERT devices were compared to those of two independent accredited laboratories in Berlin (LLBB) and Potsdam (AgroLab).

Samples were prepared using a matrix of canal water spiked with secondary effluent from the Ruhleben WWTP in Berlin. To create a well-controlled water matrix, 50 L of water were collected from the Spree Canal in Berlin (GPS: 52°31’13.3"N 13°23’44.7"E) via manual grab sampling and then filtered using a 0.45 µm filter at the beginning of the study, to remove all bacteria. Manual grab sample of WWTP secondary effluent were collected daily, filtered using a 5 µm filter to remove particle-bound bacteria, and used for spiking the canal water filtrate. Measurements targeted a single target concetration level per day, the dilution protocol targeting seven concentrations, separated by 0.4 Log_10_ unit steps on a logarithmic scale: 50,125, 313, 781, 1953, 4883, 12207 MPN/100 mL (spiking procedures assumed a baseline WWTP effluent concentration of 30000 MPN/100 mL, based on historical data).

Sample preparation occurred on-site at the WWTP, with equipment disinfected one day prior to conducting the experiment using ethanol (>70%). A total of 6.5 L sample volume was prepared for each target concentration and homogenized by vigorous turbulent mixing in a 20 L canister which was previously flushed with canal filtrate.

For the ALERT Lab metrology study, 24 aliquots of 150 mL were prepared each day for laboratory analyses (12 for each laboratory). Due to the high level of agreement observed between laboratories, only one laboratory (AgroLab) was subsequently used for the ALERT System V2 metrology study, and the number of laboratory aliquots was reduced from 12 to 8. All laboratory samples were stored in coolers for preservation during transport and storage. They arrived at the laboratories each day at noon and were processed the same afternoon within acceptable holding times.

For the ALERT Lab, aliquots of 25 mL of the homogenized sample were prepared in a beaker and loaded into each of the instrument’s six measurement vials, using a sterile syringe. For the ALERT System V2, a 100 mL beaker containing 60 mL of the sample was provided to each of the seven independent sampling inlets. The built-in temperature sensor of the ALERT System V2 was immersed in the sample beaker during sampling, to control for temperature effects of the water matrix. Sampling commands were then sent remotely from a cell phone.

### 2.3 ALERT field-deployment case studies

To demonstrate the broad applicability of ALERT technology across diverse monitoring contexts, three distinct field deployment case studies were carried out in varied environments (urban, peri-urban, and rural), across different water matrices (a canal and two major rivers), in three countries (Germany, France, and the United Kingdom). These deployments employed the different ALERT analyzers available (ALERT Lab, ALERT System V2, and ALERT One) and were operated by a range of users, including academic researchers, wastewater utility professionals, and citizen scientists.

#### 2.3.1 Berlin Spree Canal side-by-side time series

The Spree Canal in Berlin (Germany) was historically lined with public bathhouses, but is now impacted by combined sewer overflows. The canal is old arm of the river Spree and is classified as a “highly modified water body” according to the European Water Framework directive. It was a focus of multiple initiatives aimed at reopening recreational bathing sites. The ALERT deployment in the Spree Canal involved high-frequency water quality monitoring using the portable ALERT Lab analyzer in parallel with laboratory testing to evaluate the capacity of ALERT technology to provide accurate early warning in case of short-term contamination events.

Side-by-side sampling of surface waters was conducted in the Spree Canal (Berlin, Germany) on 51 days spread over a 13-week period ranging from June 17 to September 9, 2020, to evaluate the capabilities of the ALERT Lab instrument to reliably detect contamination peaks in surface waters, and compare data obtained by the ALERT technology to laboratory reference methods.

To this end, two daily manual grab samples (labeled Sample 1 and Sample 2) were collected between 9 and 10 a.m., resulting in a total of 102 samples, which were divided into four aliquots each: two were analyzed on-site in duplicate using the ALERT Lab, while the remaining two were placed in sterile bottles for duplicate analysis by an approved laboratory using the MPN method (ISO, 1998).

To assess the impact of sample preservation and delayed analysis on measured water quality, an additional aliquot from Sample 2 was re-analyzed using the ALERT Lab after undergoing the same cold storage duration as the lab samples. It was analyzed in duplicate at 3:15 p.m using the ALERT Lab, corresponding to the approximate start time of the incubation for laboratory analysis, which varied between 1:30 and 5:00 p.m.

In total, the Berlin Spree Canal campaign generated 306 ALERT measurements and 204 MPN measurements. All ALERT analyses were conducted on raw (unfiltered) samples, thus reflecting comprehensive *E. coli* concentrations.

#### 2.3.2 Seine River at Ablon-sur-Seine side-by-side time series

An *in-situ* ALERT System V2 autonomous analyzer was installed during the summer of 2021 in the Seine river at Ablon-sur-Seine (France), at a monitoring location (GPS coordinates: 48.719047, 2.415327) located upstream of Paris and downstream from the city of Orly—a major suburban hub that hosts France’s second-largest airport. The area is characterized by a hybrid sewage infrastructure comprising both combined and separate networks, which contribute to episodic sewage pollution during rainfall. The Ablon-sur-Seine ALERT deployment aimed to produce high-frequency, automated *E. coli* concentration data, enabling the characterization of water quality dynamics under both storm-driven and dry weather conditions.

The analyzer was installed directly in the river, mounted to the concrete embankment using a rail and bracket system, and programmed to perform fully automated daily sampling. Automated sampling and *E. coli* measurements were triggered daily for two months (June 1^st^ – July 28^th^, 2021). Weekly samples were manually collected by Paris region wastewater authority (SIAAP) personnel at the same location and analyzed in an approved laboratory by the MPN method (ISO, 1998). ALERT System V2 measurements were triggered simultaneously with every laboratory sample.

In addition to routine daily measurements, intensified sampling was performed during three targeted events: two rainfall episodes, during which the analyzer’s seven-cartridge capacity was used to obtain measurements over a 48-hour period; and one dry-weather episode during which seven measurements were conducted over a 24-hour span. These high-frequency sequences enabled analysis of the return-to-baseline time for E. coli concentrations following CSO-induced pollution, as well as characterization of intra-day variability under dry-weather conditions.

Over the course of the Ablon-sur-Seine study, a total of 62 ALERT measurements and 9 laboratory MPN measurements were collected. The ALERT System V2 was configured to analyze raw samples without 5 µm pre-filtration, therefore providing comprehensive *E. coli* measurements.

#### 2.3.3 Thames River at Henley-on-Thames citizen-science time series

A community-driven field monitoring initiative on the River Thames at Henley-on-Thames—a scenic location known for the Henley Royal Regatta but significantly impacted by CSOs. The Henley-on-Thames monitoring campaign began in January 2025 and remains ongoing. Conducted entirely by a local citizen-science volunteer group, HoT (Henley-on-Thames) Water, the study aims to deliver high-frequency data for river users and better understand the effects of infrastructure failures on water quality. The HoT initiative received the Fluidion Water Safety Excellence Certification label and adheres to its quality chart (https://fluidion.com/customer-resources/certification-program).

To date, 72 samples have been collected from four nearby river locations, with two sites sampled per day. Each sample was analyzed for comprehensive and planktonic *E. coli* counts using multiple handheld ALERT One analyzers, to obtain long-term high-frequency water quality trends at the historic location of the Royal Henley Regatta and quantify the presence of aggregate-bound *E. coli* bacteria.

### 2.4 Evaluation of point-of-use water treatment solutions

This part of the study applied applied ALERT field instrumentation to assess the microbiological abatement efficacy at progressively decreasing concentrations of a common point-of-use potabilization treatment designed for LMICs, so as to identify the minimum dosage requirement and optimize cost effectiveness without compromising safety in resource-constrained settings.

Grab samples were collected over a two-month period (February 25 – April 24, 2025) in the Seine river at Alfortville (GPS coordinates 48.813444, 2.410961), an urban site heavily polluted by both upstream river pollution and nearby discharge of effluent from the Valenton WWTP, a major plant having a treatment capacity of 600,000 m^3^ / day.

Samples were treated using P&G Purifier of Water Packet (Procter & Gamble), a widely-used flocculation-disinfection treatment verified by WHO to provide point-of-use comprehensive water treatment (WHO, 2025) using standard manufacturer-recommended procedures: the P&G powder was added and well-mixed into 10 L of river water, which was then allowed to flocculate (5 minutes), filtered using cotton cloth, and then allowed to rest for and additional 20 minutes.

*E. coli* concentrations were evaluated at different stages of the treatment process: 1. as sampled (non-treated, non-filtered – NTNF); 2. following addition of the P&G disinfectant and flocculation (treated, non-filtered – TNF); and 3. following filtration using cotton cloth and 20 minutes rest (treated, filtered – TF). Sodium thiosulfate was added after the extraction of the aliquot for analysis to quench the disinfection reaction.

The minimal treatment dose required for producing microbiologically safe water on-site was determined by performing 28 separate experiments using progressively lower concentrations of P&G powder, from the manufacturer-recommended dose (100% – 4 samples), down to one-half (50% – 2 samples), one-quarter (25% – 5 samples), and one-eighth strength (12.5% – 17 samples). The relative treatment efficacy for inactivating planktonic and, respectively, aggregate-bound *E. coli* bacteria was evaluated by analyzing samples raw (for comprehensive *E. coli* counts) and following 5µm filtration (for planktonic *E. coli* counts). *E. coli* concentration measurements were performed using both ALERT technology (ALERT Lab and ALERT One instruments) and the laboratory MPN method IDEXX Colilert Quantitray-2000, following standardized protocols (ISO, 2012).

In total, the study generated 168 ALERT and 168 MPN measurements.

### 2.5 Microscopic *E. coli* observations

#### 2.5.1 *E. coli* immunostaining procedure

##### Phosphate-buffered saline (PBS) solution preparation

1 tablet of PBS pH 7.4 (Roth, REF: 1112) was dissolved in 1 L of autoclave-sterilized deionized water (Millipore Elix® Essential 3 Water Purification System).

##### Fixation, filtration and permeabilization

A volume of 20 mL of the sample was mixed with 60 mL of 4 % paraformaldehyde (ThermoFisher, Ref.: 30525-89-4) in PBS and kept 30 min at ambient temperature. The resulting mixture was then filtered in 10 mL inclements, using a hydrophilic black Nucleopore track-etched polycarbonate membrane filter (47 mm diameter, 0.2 μm pore size, Whatman, Ref.: 111156) coupled to a laboratory vacuum pump (KNF). The filter was then rinsed with 10 mL of PBS solution, followed by 10 mL of PBS allowed to contact the membrane for 4 min before vacuum filtration and a final 10 mL PBS rinse. Volumes of 40 µL of sheep serum (ThermoFisher, Ref.: 16070096) and 0.8 µL of Triton X-100 (ThermoFisher, Ref.: 9002-93-1) were mixed with PBS solution to obtain a final 400 µL volume of buffer, which was pipetted over the filter and subsequently left 1 h at ambient temperature. The membrane was then rinsed with 20 mL of PBS solution.

##### Antibody staining

Volumes of 40 µL of sheep serum, 0.8 µL of Triton X-100 and 20 µL of *E. coli* serotype 0/K Antibody (ThermoFisher, Ref.: PA1-73029) were mixed with PBS solution to obtain a final 400 µL volume of staining solution, which was pipetted over the filter, ensuring complete coverage. The filtration apparatus was then stored in the dark for 2 h, then the membrane was rinsed with 20 mL of PBS solution.

##### DAPI staining

A volume of 0.6 µL of DAPI (ThermoFisher, Ref.: 0062248) was mixed with PBS solution to obtain a 400 µL volume staining solution, which was pipetted over the filter, ensuring complete filter coverage. The filtration apparatus was then stored in the dark for 8 min, then the membrane was rinsed with 20 mL of PBS solution and air dried.

#### 2.5.2 Epifluorescence microscopy of immunostained bacteria

##### Microscopy setup

A Leica DMI 3000M inverted fluorescence microscope was used for all imaging. It was outfitted with an oil immersion objective (Leica N PLAN 50x/0,90 OIL), a fiber-coupled mercury lamp, and the A and I3 fluorescence dichroic filter cubes that allowed differential imaging of the DAPI and FITC fluorophores.

##### Sample preparation

The Nucleopore filter containing the dyed sample was carefully removed from the filtration apparatus using round-tipped sterilized metal tweezers. It was then cut in approximately 1 cm^2^ portions using scissors. A portion of the filter was then placed over a glass slide wet with anti-fade solution (SlowFade, ThermoFisher, Ref.: S36968) and secured to it using a small drop of N-type immersion oil. Additional drops of N-type immersion oil were placed over the filter and on the objective, respectively. The glass slide was then placed on the microscope stage. Imaging: Image recording was performed using a FHD V2.0 C-mount color camera, and images were saved to an SD memory card for further processing (background subtraction, noise reduction, contrast enhancement, scale display) on a computer, using the Gimp (https://www.gimp.org/) and Fiji ImageJ (https://imagej.net/software/fiji/) packages.

## 3 Results

The results and calculations presented in this study rely on the underlying assumption that microbiological measurements follow a log-normal distribution, and most statistical calculations will therefore use Log_10_-transformed values. Across the different parts of this study, a total of 705 samples were analyzed using ALERT technology implemented in three different types of ALERT rapid microbiology analyzers, and 471 side-by-side samples were analyzed using the MPN laboratory method, without including samples used in the independent testing performed by KWR Laboratories for the WHO drinking water verification protocol. The amount of side-by-side data collected makes this study, to our knowledge, the most thorough evaluation reported to date of any rapid microbiology technology.

### 3.1 ALERT metrology study

Figure 2 displays graphically the results of the side-by-side metrology studies for ALERT Lab and ALERT System V2. Excellent linearity is observed across the full range of concentrations for both the ALERT Lab (R² = 0.9962) and the ALERT System V2 (R² = 0.9941). A constant offset of +0.31 Log_10_ units was recorded for the ALERT Lab, while for the ALERT System V2, the recorded offset was -0.28 Log_10_ units. The insets display actual side-by-side datapoints from the different spiked standard solutions used, to give an idea of their sample-to-sample statistical dispersion. In the left panel, ALERT Lab measurements 1-3 were paired with aliquots 1-3 from AGRO Lab, while ALERT Lab measurements 4-6 were paired with MPN measurements 1-3 from LLBB Lab. In the right panel, ALERT System V2 measurements 1-7 were paired with MPN measurements 1-7 from AGRO Lab. The colors in the insets identify the individual spiked standard solution from which multiple aliquots were analyzed, side by side, by ALERT and the laboratory.

**Figure 2.**
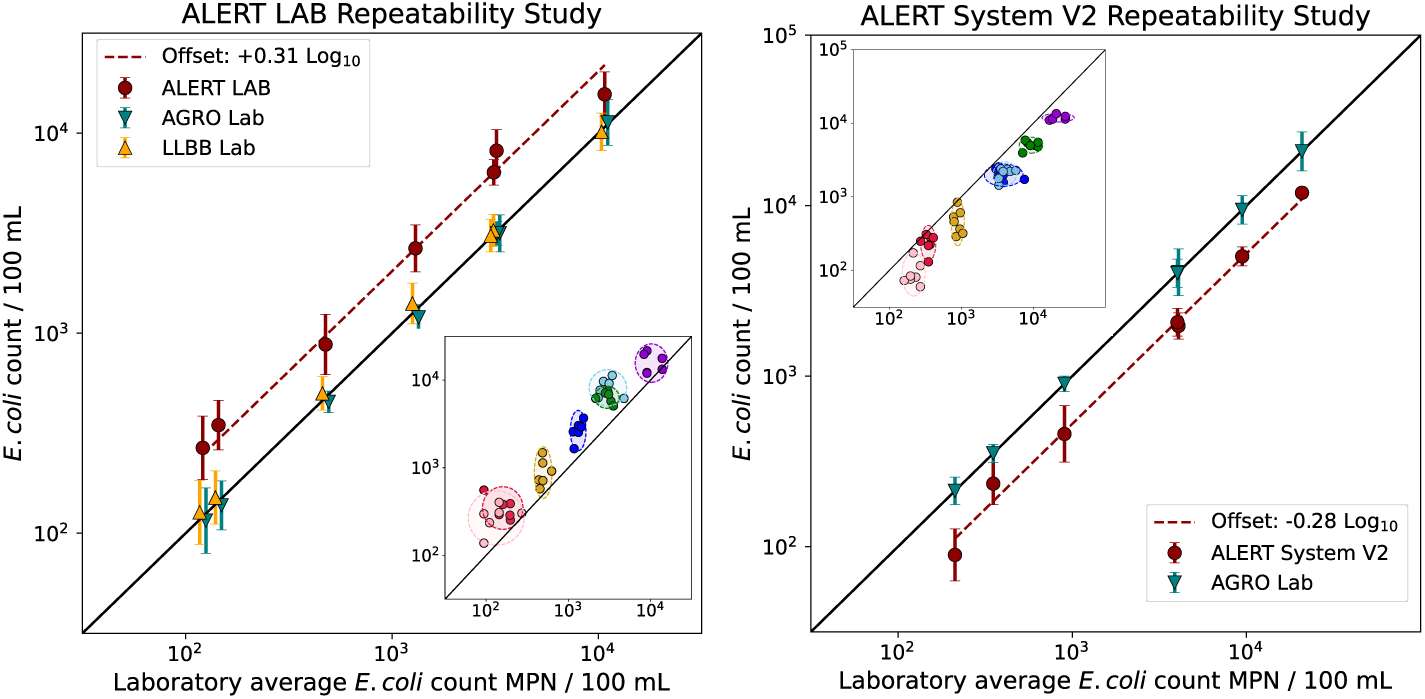
ALERT vs. laboratory MPN comparison (at left: ALERT Lab, at right: ALERT System V2). The horizontal axis represents the average of all laboratory values, while the vertical axis displays the average value per concentration and per method, with vertical error bars representing the corresponding standard deviation (SD). Data points were slightly shifted horizontally for clarity. Insets: side-by-side representations, with individual spiked standard solutions identified by the same color.

Precision, as measured by the standard deviation (SD) of the repeat measurements on each standard solution, was similar to that of the standard laboratories for the ALERT Lab (Table 2). The SD ranged from a maximum of 0.16 Log_10_ units at the lower concentration end, to 0.06-0.11 Log_10_ units at higher concentrations.

**Table 2.**
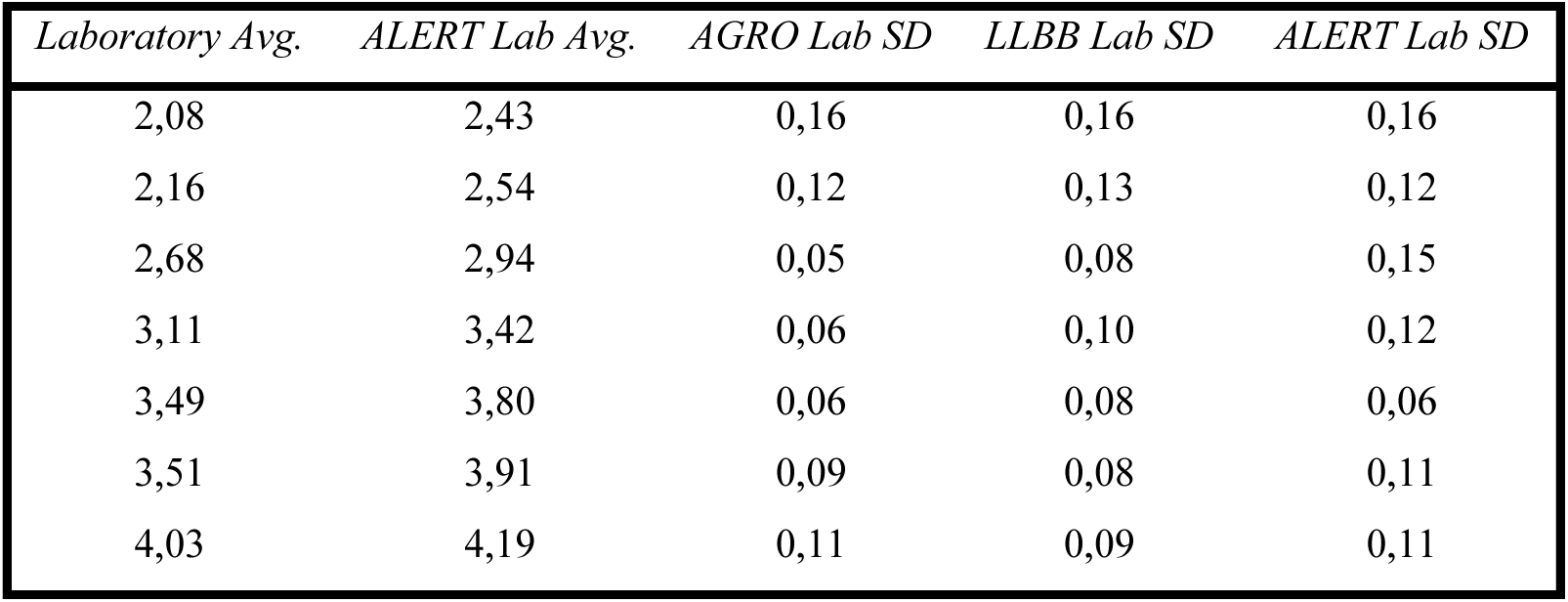
ALERT Lab vs. MPN metrology comparison (all values are presented in Log_10_ units).

| *Laboratory Avg.* | *ALERT Lab Avg.* | *AGRO Lab SD* | *LLBB Lab SD* | *ALERT Lab SD* |
| --- | --- | --- | --- | --- |
| 2,08 | 2,43 | 0,16 | 0,16 | 0,16 |
| 2,16 | 2,54 | 0,12 | 0,13 | 0,12 |
| 2,68 | 2,94 | 0,05 | 0,08 | 0,15 |
| 3,11 | 3,42 | 0,06 | 0,10 | 0,12 |
| 3,49 | 3,80 | 0,06 | 0,08 | 0,06 |
| 3,51 | 3,91 | 0,09 | 0,08 | 0,11 |
| 4,03 | 4,19 | 0,11 | 0,09 | 0,11 |

For the ALERT System V2 (Table 3), measured precision was similarly comparable that of the standard laboratory across the full concentration range, with notable improvement compared to the laboratory MPN method at the higher concentrations. An important design feature of the ALERT System V2 resides in the automation of all sampling in sterile disposable measurement cartridges, which completely eliminates potential human error and contamination that could potentially occur during manual sampling and sample preparation operations, while ensuring that performance metrics evaluated in the laboratory translate to field deployments.

**Table 3.** ALERT System V2 vs. MPN metrology comparison (all values are presented in Log_10_ units)

| *AGRO Lab Avg.* | *ALERT System V2 Avg.* | *AGRO Lab SD* | *ALERT System V2 SD* |
| --- | --- | --- | --- |
| 2,33 | 1,95 | 0,08 | 0,15 |
| 2,55 | 2,37 | 0,05 | 0,12 |
| 2,96 | 2,66 | 0,05 | 0,17 |
| 3,60 | 3,32 | 0,08 | 0,08 |
| 3,61 | 3,29 | 0,14 | 0,08 |
| 3,98 | 3,70 | 0,08 | 0,06 |
| 4,32 | 4,08 | 0,11 | 0,03 |

### 3.2 ALERT field deployment case studies

The results from the Berlin Spree Canal study are presented in Figure 3, including time-series plots and side-by-side comparisons of on-site ALERT versus laboratory MPN measurements, as well as ALERT measurements following cold storage versus MPN results. Duplicate MPN readings were also plotted side by side to comparatively illustrate the internal reproducibility of the reference method.

**Figure 3.**
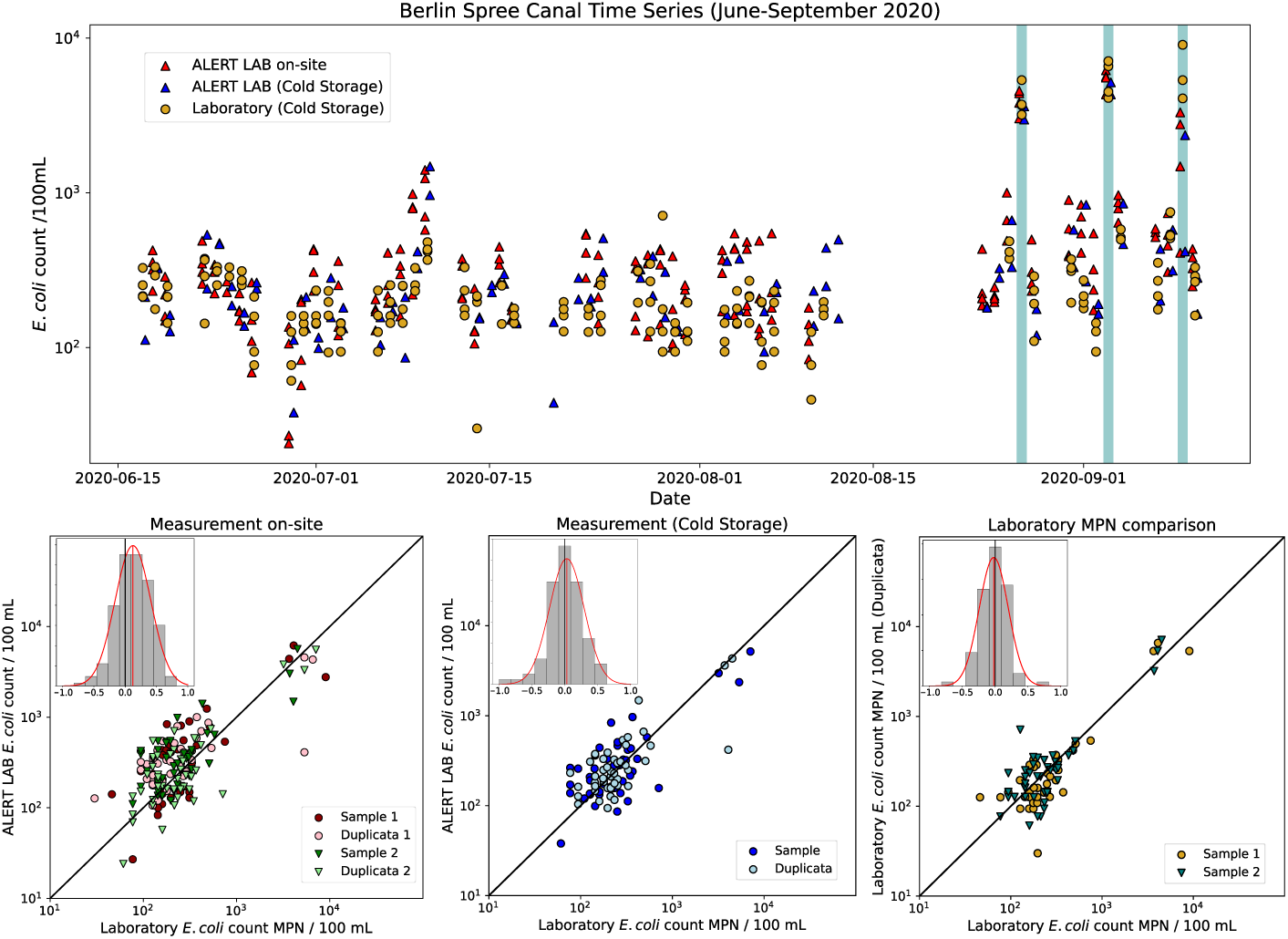
Berlin Spree Canal study results. Top panel: time-series data, with three consecutive STP incidents at the end of the monitoring period highlighted using blue vertical bars. Bottom panels represent side-by-side data. Left: ALERT on-site measurements vs. laboratory MPN; Middle: ALERT measurement following cold storage vs. laboratory MPN; Right: Comparison of duplicata laboratory MPN measurements. Bottom panel insets present the histograms of respective mesurement differences, with overlaid gaussian plots of equivalent mean and standard deviation.

Overall, the Spree Canal water during the first two months of the 2020 monitoring season (June 15– August 15) was classified as good quality, according to the European bathing water directive (EU, 2006) (ALERT comprehensive geometric mean (GM): 2.38 Log_10_ (equivalent to 240 *E. coli*/100 mL), SD: 0.28 Log_10_, upper 95^th^ percentile: 2.85 Log_10_ (or 704 *E. coli*/100 mL)). However, toward the end of the monitoring period, three consecutive short-term *E. coli* contamination events were detected, as illustrated in the Figure 3 time-series graph. When assessed over the full study duration, comprehensive GM was 2.51 Log_10_ (322 *E. coli* / 100 mL), with a SD of 0.39 Log_10_ and the upper 90^th^ percentile at 3.01 Log_10_ (1013 *E. coli* / 100 mL), implying insufficient quality according to the European bathing water directive (EU, 2006), unless the bathing site were closed during the short-term contamination events.

All three contamination events were accurately identified and classified by both the ALERT system and laboratory-based MPN analyses. Notably, ALERT provided quantitative results and issued early warnings within 8 hours of sampling, while laboratory results became available only several days later.

The side-by-side analysis enables quantification of the agreement between the ALERT system and the laboratory MPN method under real-world field conditions, using raw environmental samples that may contain bacterial aggregates—unlike the metrology study, where the spiking matrix was pre-filtered. A slight positive offset was observed, with ALERT measurements obtained on-site immediately after sampling showing a mean overestimation of 0.12 Log_10_ units relative to the laboratory MPN results. The distribution of differences between the two methods approximates a Gaussian profile, with a SD of 0.29 Log_10_ units (Figure 3, inset, bottom-left panel). The calculated Pearson correlation coefficient between the ALERT and MPN results is *r=*0.75, and the index of agreement *IA*=0.85, calculated using the alternative methods calculator tool provided by the USEPA [USEPA, 2021].

It is instructive to compare the agreement of MPN and ALERT counts in terms of GM value and statistical threshold value (STV), regulatory risk assessment parameters whose exceedances may trigger actions such as temporary or permanent closures of a recreational site. Agreement matrices were computed for the GM=900 (90^th^ percentile) used to classify inland water as “sufficient” under the European bathing water directive (EU, 2006), and for the STV=1800 thershold applied in certain European countries for immediate open/close decisions. The measured agreement rate was 97.1% (GM) and, respectively, 98.9% (STV), indicating excellent agreement between the MPN and ALERT methods on regulatory water quality classification.

**Table 4.**
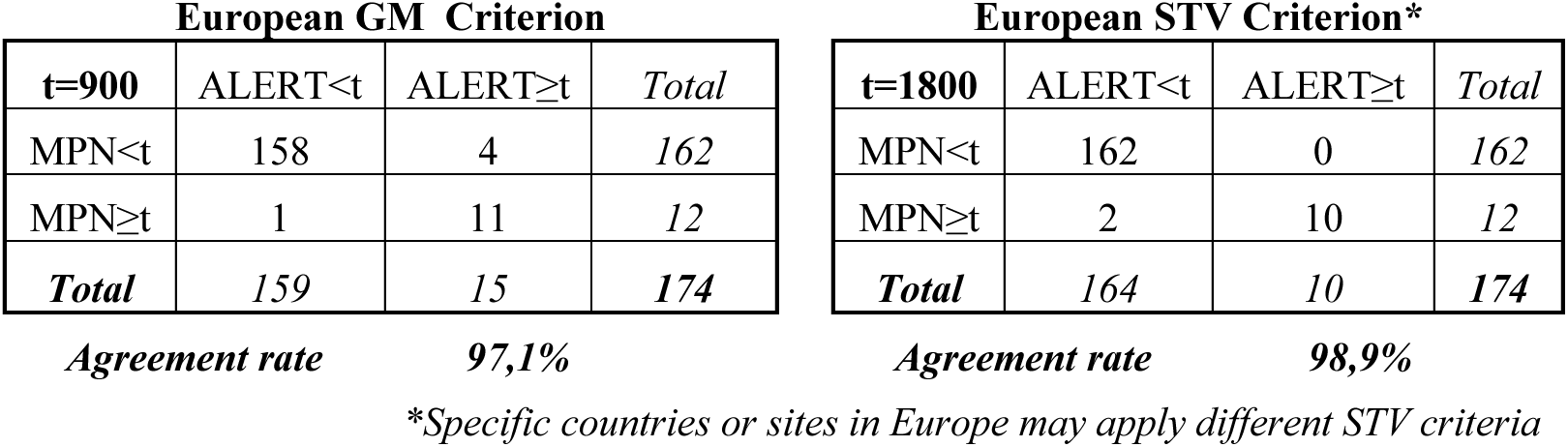
Agreement matrices for E. coli concentration results obtained by the MPN and ALERT methods on raw samples, in terms of the STV and GM threshold criteria recommended in Europe (right two panels)

| European GM Criterion |  |  |  | European STV Criterion* |  |  |  |
| --- | --- | --- | --- | --- | --- | --- | --- |
| t=900 | ALERT<t | ALERT≥t | Total | t=1800 | ALERT<t | ALERT≥t | Total |
| MPN<t | 158 | 4 | 162 | MPN<t | 162 | 0 | 162 |
| MPN≥t | 1 | 11 | 12 | MPN≥t | 2 | 10 | 12 |
| **Total** | **159** | **15** | **174** | **Total** | **164** | **10** | **174** |
| *Agreement rate* |  | **97,1%** |  | *Agreement rate* |  | **98,9%** |  |

*\*Specific countries or sites in Europe may apply different STV criteria*

When analyzing the samples after cold storage, the positive offset of ALERT versus MPN is greatly reduced (0.03 Log_10_ units), becoming negligible compared to the stardard laboratory measurement uncertainty, while the distribution of differences shows a slightly reduced SD of 0.26 Log_10_ units.

A field precision estimate can be obtained by applying the distribution-of-differences analysis on duplicata ALERT and MPN laboratory measurements and thereby determine the internal reproducibility of the two methods. The distribution of ALERT duplicata measurement differences provided a mean of 0.04 Log_10_ units and a SD of 0.27 Log_10_ units, while that of MPN duplicata measurement differences provided a mean of 0.02 Log_10_ units and a SD of 0.22 Log_10_ units. From these values we can infer the respective precision of the ALERT and MPN laboratory measurements in field conditions and using real raw samples, by dividing the SD of the differences by √2. We obtain a precision of 0.19 Log_10_ units (ALERT) and 0.16 Log_10_ units (MPN), corresponding to a baseline geometric mean of 2.38 Log_10_.

Figure 4 (top panel) displays the Ablon-sur-Seine time series results along with rainfall data recorded hourly at Orly (for better visibility, a six-hour rolling average was applied to the rainfall data). Several rain events were recorded during the study period, leading to increased *E. coli* concentrations. Two storm events (June 4-6 and 17-19, 2021) were closely monitored through high-frequency ALERT sampling campaigns (Figure 4, bottom left and center panels).

**Figure 4.**
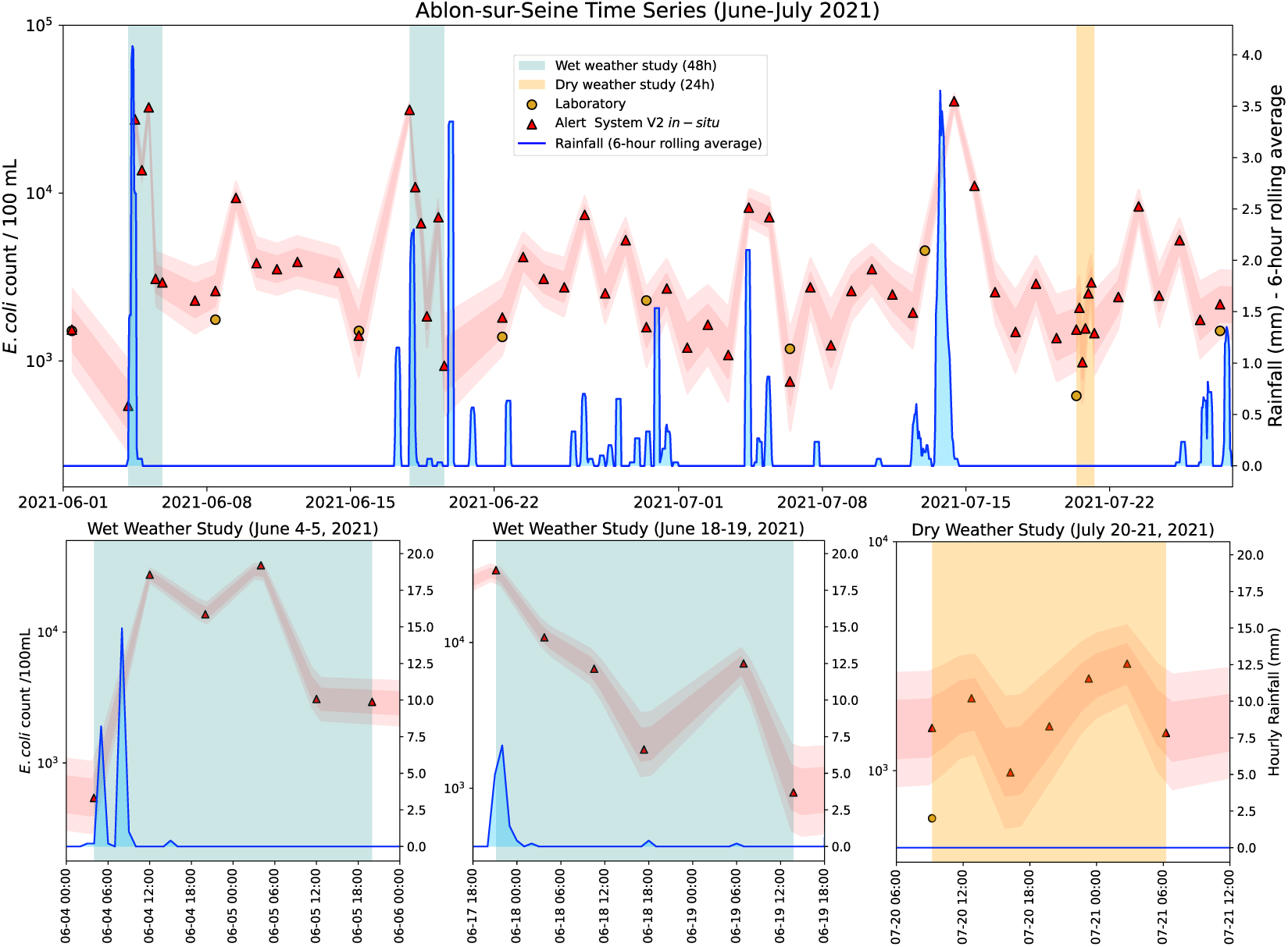
Results from the Ablon-sur-Seine monitoring study. Top panel: Time-series plot showing E. coli concentrations measured by the ALERT System V2 and the laboratory MPN method. Red bands around ALERT data represent the SD and 95% confidence intervals, as derived from Table 3. Vertical blue and yellow shading indicate high-frequency sampling periods during storm events and dry weather, respectively. Bottom panels: Enlarged views of the high-frequency sampling campaigns, highlighting temporal dynamics under different hydrological conditions.

During the June 4 storm event (Figure 4, bottom left panel), six ALERT measurements were taken at 8-hour intervals, beginning at 4:00 a.m., coinciding with the onset of rainfall (*E. coli* concentration at start: 541 CFU/100 mL). Rainfall intensity peaked at 5:00 a.m. (8.2 mm/h) and again at 8:00 a.m. (14.9 mm/h). Two distinct *E. coli* concentration peaks were recorded: the first at 12:00 p.m. on June 4 (27,459 CFU/100 mL) and the second at 4:00 a.m. on June 5 (32,408 CFU/100 mL). Following these peaks, concentrations declined to 3,082 and 2,932 CFU/100 mL at 12:00 p.m. and 8:00 p.m. on June 5, respectively—values consistent with the site’s dry-weather baseline.

During the June 17 storm event (Figure 4, bottom middle panel), two rainfall intensity peaks were recorded—one at 5:00 a.m. (4.7 mm/h) and another between 9:00 p.m. and 10:00 p.m. (4.9 mm/h and 6.9 mm/h, respectively). Seven ALERT System V2 measurements were conducted at 7-hour intervals, beginning at the onset of the second rainfall peak. One data point (12:00 a.m. on June 19) was not recorded due to a measurement cartridge malfunction.

The first measurement, taken at 9:09 p.m. on June 17, captured the peak *E. coli* concentration of the event (31,352 CFU/100 mL). This was followed by a gradual decline over the next 40 hours, with a secondary peak of 7,174 CFU/100 mL observed at 6:57 a.m. on June 19. By 1:45 p.m. that same day, concentrations had returned to baseline levels (935 CFU/100 mL). Since no samples were collected immediately following the first rainfall peak, it is possible that *E. coli* levels may have peaked earlier and exceeded the recorded maximum.

A third major contamination peak, driven by rainfall, was recorded at 10:11 a.m. on July 14, with *E. coli* levels reaching 35,208 CFU/100 mL following 40.9 mm of precipitation over the preceding 24 hours. Although the dynamics of the event were not captured through high-frequency sampling, valuable insights were obtained from subsequent daily measurements. Within 24 hours, *E. coli* concentrations had dropped to 11,038 CFU/100 mL, and returned to baseline dry-weather levels (2,568 CFU/100 mL) by 10:04 a.m. on July 16—48 hours after the peak.

The July 20–21 high-resolution monitoring campaign (Figure 3, bottom right panel) focused on a dry-weather period following a full week without rainfall. Seven ALERT measurements were taken at 3.5-hour intervals, beginning on July 20 at 9:15 a.m. The results revealed two distinct daily peaks in *E. coli* concentrations: 2,071 CFU/100 mL at 12:45 p.m. on July 20, and 2,932 CFU/100 mL at 2:45 a.m. on July 21. The lowest concentration during the period—983 CFU/100 mL—was recorded at 4:15 p.m. on July 20.

Based on insights from the June 4 storm event, which indicated a 4–7 hour travel time for pollution to reach Ablon-sur-Seine from Orly, the July peaks would correspond to upstream discharges occurring between 5:45–8:45 a.m. and 7:45–10:45 p.m.—coinciding with peak domestic water use periods on working days. These fluctuations could indicate dry-weather discharges originating from Orly, either through sewage mis-connections into stormwater drains in areas served by separate sewer systems, or from live-aboard boats discharging untreated wastewater directly into the river.

Analysis of the side-by-side ALERT and laboratory MPN data reveals strong agreement, despite the limited dataset of only nine paired measurements. The offset between the two methods was minimal (0.06 Log_10_ units), and the differences were tightly distributed, with a SD of only 0.19 Log_10_ units. Over the monitored period, the laboratory MPN GM was 3.20 Log_10_ units (1,587 *E. coli* / 100 mL) and the ALERT GM was 3.50 Log_10_ units (3,155 *E. coli* / 100 mL), both agreeing on the classification of insufficient water quality, according to the European bathing water directive (EU, 2006).

A slightly different picture emerges when comparing the respective SD, 0.23 Log_10_ units for the laboratory and 0.40 Log_10_ units for ALERT, which indicate much higher variability of water quality when measured at daily frequency (using ALERT technology) than weekly (using the laboratory). Indeed, it can be seen from Figure 4 that all weekly laboratory samples were collected during dry-weather conditions and did not capture the fluctuations due to STP events that could be observed through higher frequency ALERT measurements.

The monitoring data from the Henley-on-Thames case study are presented in Figure 5, alongside rainfall records and reported CSO events from the Wargrave wastewater treatment plant (WWTP), located upstream of the sampling sites. Overall, the planktonic *E. coli* GM for this study was 2.06 Log_10_ units (114 *E. coli* / 100 mL), SD 0.39 Log_10_ units, and the upper 95th percentile at 2.69 Log_10_ units (485 *E. coli* / 100 mL), indicating excellent water quality classification according to European bathing water directive (EU, 2006). The comprehensive *E. coli* GM was 2.68 Log_10_ units (478 *E. coli* / 100 mL), with a SD of 0.48 Log_10_ units and an upper 90th percentile of 3.24 Log_10_ units (1,786 *E. coli* / 100 mL), indicating insufficient water quality classification.

**Figure 5.**
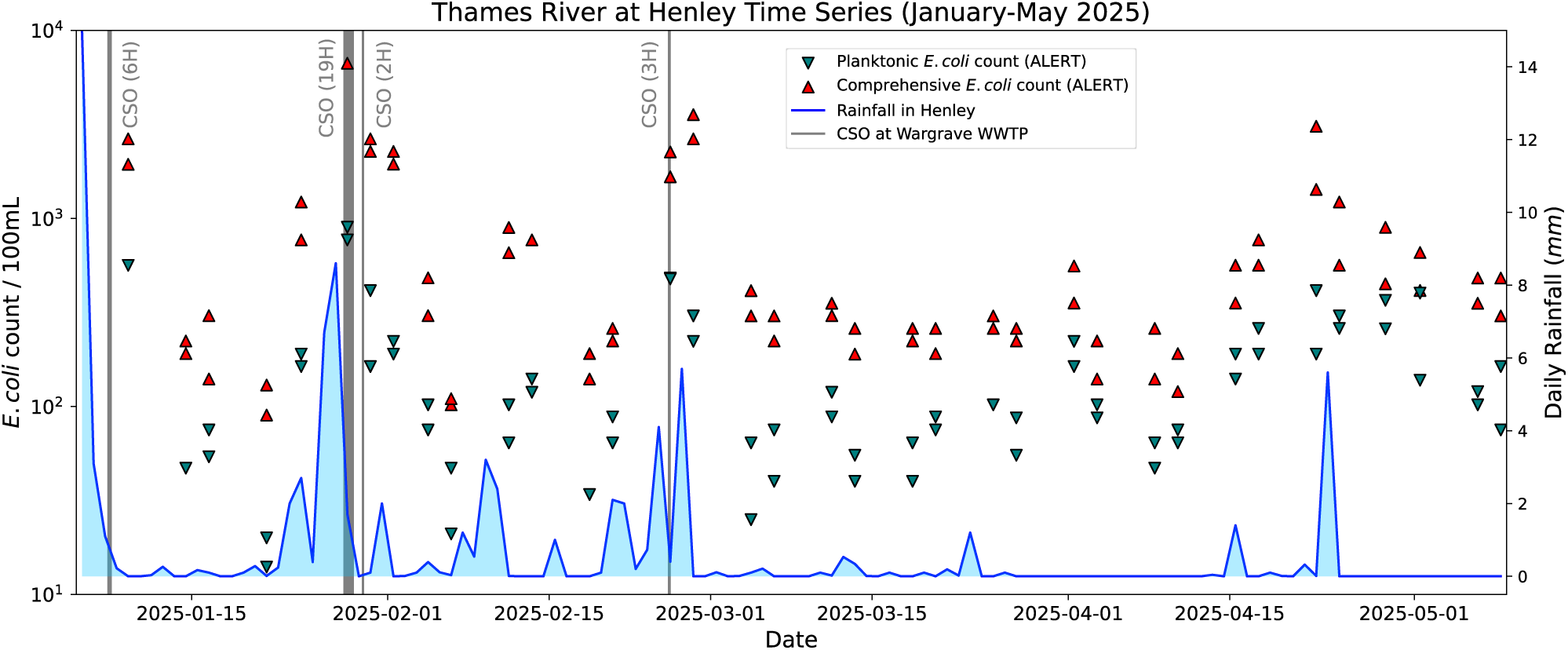
Thames River study (Henley-on-Thames). Comprehensive and planktonic E. coli counts are presented along with daily rainfall. Two nearby locations were sampled on each day. Reported CSO events are represented as vertical bars, their width corresponding to the event duration.

A strong correlation is observed between CSO discharges and significant spikes in *E. coli* concentrations. The highest recorded values—comprehensive *E. coli* counts of 6,689 *E. coli* / 100 mL at both Marsh Lock and the Rowing Museum—were measured on January 28, 2025, coinciding with the largest CSO event during the study period. Notably, the planktonic *E. coli* concentrations in these samples were much lower (898 and 769 *E. coli* / 100 mL, respectively), reinforcing the hypothesis that fecal particles, which dominate in untreated discharges, carry a disproportionately high bacterial load. When averaged across all the samples analyzed in the study, the fraction of planktonic bacteria corresponded to 27%, the remaining 73% being accounted for by aggregate-bound *E. coli*.

Even relatively short CSO events, such as the one on February 25, 2025, can have a substantial and lasting impact. Within two days, comprehensive *E. coli* concentrations reached 2,650 CFU/100 mL at Hanley Bridge and 3,566 CFU/100 mL at Fawley Meadows, while planktonic counts remained significantly lower (303 and 222 CFU/100 mL, respectively), once again highlighting the pronounced contribution of particulate fecal matter pollution.

No additional CSO discharges were reported during the extended dry period following this event, and the ratio of comprehensive to planktonic *E. coli* counts decreased over time (Figure 6), with an estimated decay time constant of 27 days — progressively dropping from a peak value of 1.07 Log_10_ units (equivalent to a factor of 11.8) on February 27 to a baseline value of 0.39 Log_10_ units (factor of 2.4).

**Figure 6.**
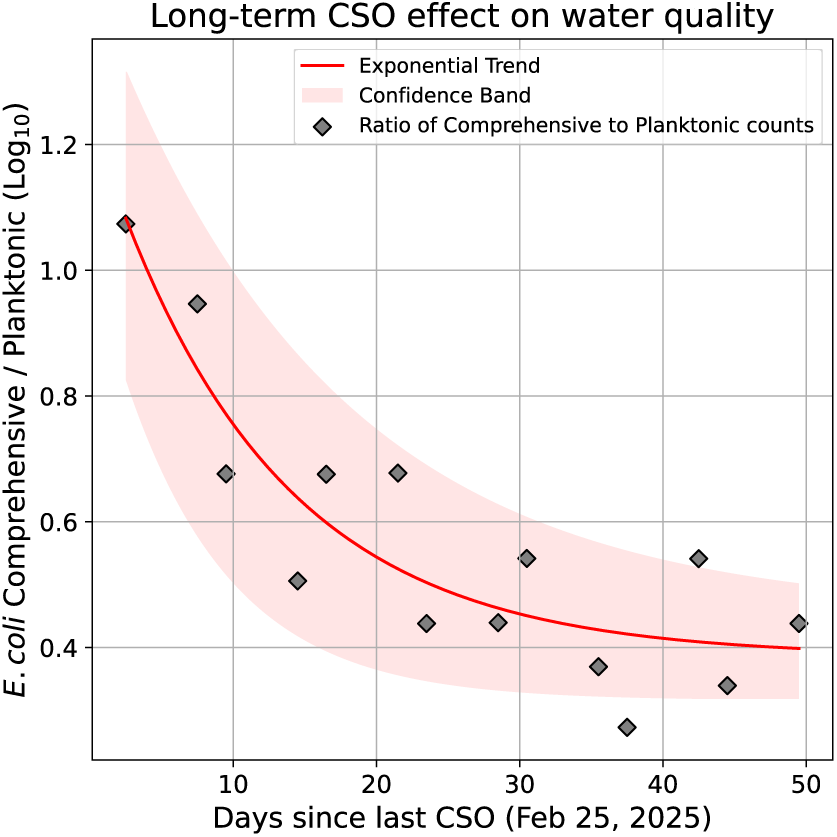
Ratio of comprehensive to planktonic E. coli counts (gray diamonds, logarithmic scale) showing a slow, gradual, decline over time during the dry period following a short-duration CSO event. Same-day samples from multiple locations were combined using the geometric mean. An exponential fit is overlaid (bright red line), with its confidence interval shown as a dim red band.

Elevated *E. coli* levels were also detected during and after rainfall events without any reported CSO activity — for example, on April 22 — suggesting either upstream CSO discharges outside the Wargrave WWTP catchment or possible gaps in CSO event reporting.

### 3.3 ALERT for LMIC-relevant applications

#### 3.3.1 Point-of-use water treatment solution evaluation

Among the samples treated at 100% P&G concentration, none showed presence of *E. coli* post-flocculation or following the full treatment. At 50% P&G concentration, samples showed single-digit positives post-flocculation, but no positives in the fully treated water. At 25%, all samples showed presence of *E. coli* post-flocculation, and occasional positives in fully treated water. Finally, at 12.5% treatment strength, all samples except two showed presence of *E. coli* post-flocculation, and all samples except four in the fully treated water.

The substantial presence of fecal aggregate-bound *E. coli* in the Seine River, as previously documented (Angelescu et al., 2024a), makes this matrix particularly well suited to evaluating the differential treatment efficacy on free-floating versus aggregate-bound bacteria.

The Seine samples collected for this study showed widely variable *E. coli* concentrations covering over two orders of magnitude. Across all the Seine river samples included in this study, the average fraction of planktonic *E. coli* bacteria was 39%, the remaining 61% corresponding to aggregate-bound *E. coli*.

Overall, the planktonic *E. coli* GM for this part of the study was at 2.94 Log_10_ units (868 *E. coli* / 100 mL), the SD at 0.53 Log_10_ units, and the upper 90th percentile at 3.62 Log_10_ units (4144 *E. coli* / 100 mL). From a bathing water perspective, these numbers indicate insufficient water quality classification at the sampling location over the period of the study, according to the European bathing water directive (EU, 2006). The comprehensive *E. coli* GM of 3.41 Log_10_ units (2,591 *E. coli* / 100 mL) further confirms the insufficient water quality classification.

Figure 7 provides new microscopic evidence for the presence of aggregate-bound *E. coli*, showing an epifluorescence micrograph of a typical aggregate collected from the Seine. The numerous *E. coli* attached to the DNA-rich particle support its fecal origin hypothesis (as opposed to, for example, *E. coli*-colonized mineral sediment particles). Until now, the response of such fecal aggregates to disinfection treatments has not been systematically documented.

**Figure 7.**
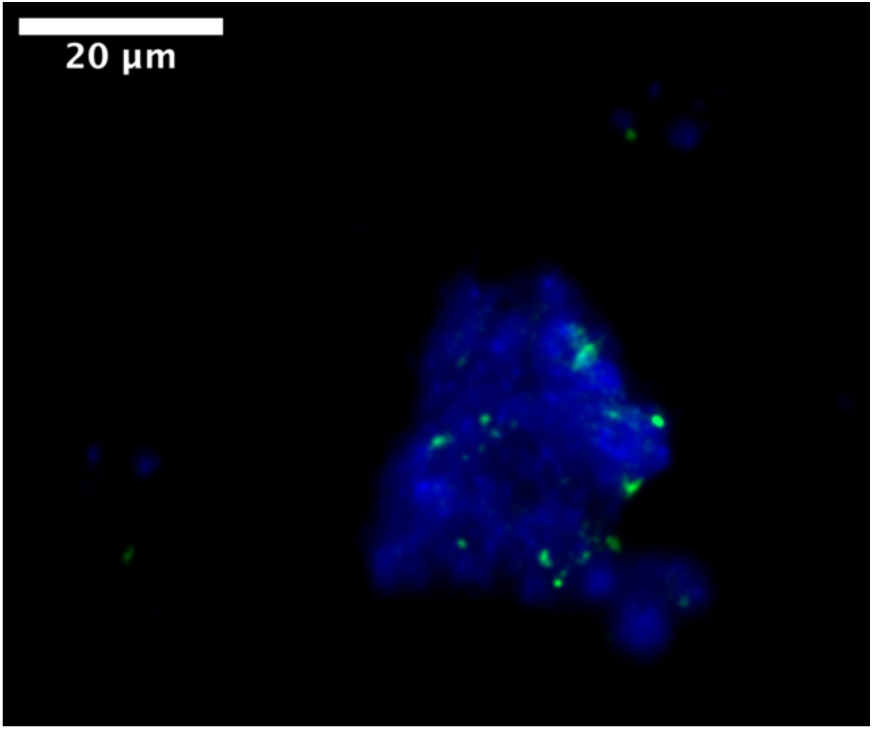
Epi-fluorescence micrograph of E. coli bacteria (green fluorescence from species-specific immunostaining) aggregated to a DNA-rich fecal particle (blue fluorescence from non-specific DNA staining)

Figure 8 presents a graphical summary of the study results at the one-eighth (12.5%) treatment strength, separating comprehensive, planktonic, and aggregate-associated *E. coli* counts (with the aggregate fraction calculated as the difference between the first two). The results clearly show that while the majority of the bacterial load resides in the aggregated fraction, the treatment is markedly more effective at reducing planktonic *E. coli* than aggregated forms.

**Figure 8.**
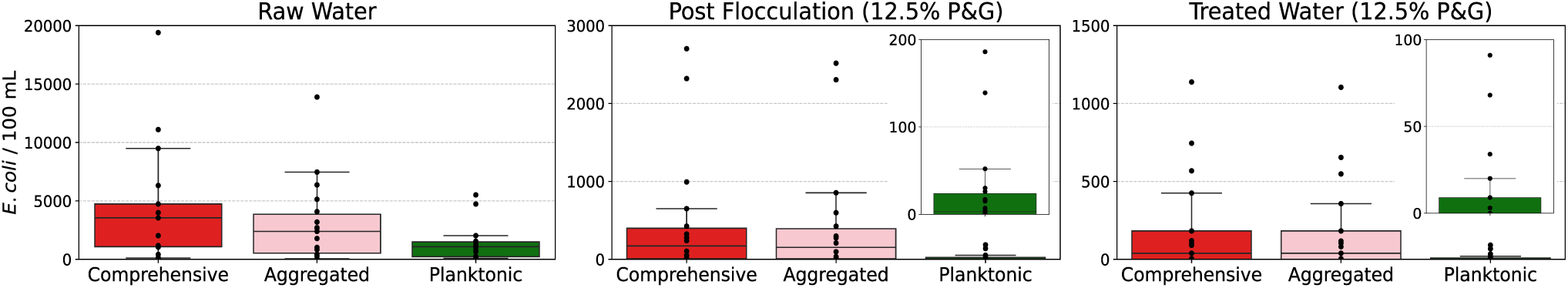
Evaluation of the P&G point-of-use water treatment solution at one-eighth strength (12.5%). The three panels illustrate E. coli concentrations across the different treatment stages: raw water (left), post-flocculation (center), and fully treated water (right). Each panel presents the distribution as bar plots for comprehensive counts (red), planktonic counts (green), and aggregated counts (pink). Box plot elements include the median (center line), interquartile range (IQR; box edges), and whiskers extending to the smallest and largest values within 1.5× IQR. For better visibility, insets are provided in the middle and right panels providing enlarged box plots of the planktonic E. coli counts.

Table 5 summarizes the calculated abatement factors for all samples testing positive at the post-flocculation and full treatment stages. Notably, the measured abatement factors for the planktonic fraction were significantly higher than those for the aggregated fraction (+0.65 Log₁₀ post-flocculation, corresponding to a 4.5-fold greater reduction; +0.59 Log₁₀ after full treatment, or 3.9-fold greater reduction). These findings suggest that *E. coli* bacteria aggregated onto fecal particles are substantially less responsive to treatment compared to their free-floating counterparts.

**Table 5.** Treatment efficacy for one-eigths (12.5%) strength P&G treatment. E. coli abatement factors are presented both post-flocculation and post-treatment, for the planktonic and aggregated E. coli fractions, as well as for the comprehensive E. coli count.

|  | Planktonic | Aggregated | Comprehensive |
| --- | --- | --- | --- |
| Post-flocculation abatement (Log<sub>10</sub>) | 1.85 | 1.20 | 1.36 |
| Full treatment abatement (Log<sub>10</sub>) | 1.94 | 1.35 | 1.52 |

#### 3.3.2 Drinking water quality evaluation

As part of UNICEF’s Rapid Water Quality Testing initiative, ALERT technology was independently tested and verified according to WHO protocols by the KWR Laboratory in the Netherlands. The testing report is publicly available (WHO-UNICEF, 2022), the results being summarized below.

A total of 60 paired samples were tested side-by-side against the IDEXX Colilert-Quantitray standard laboratory method (ISO, 2012) across four artificially spiked waters and five natural water matrices. In 87% of tests, the ALERT semi-quantitative risk class matched the value provided by the laboratory measurement. Whenever there was a discrepancy, ALERT tended to overestimate rather than underestimate risk—supporting its use as a conservative drinking water safety tool.

When used as a presence/absence test, ALERT correctly classified 93% of samples using a 1 CFU/100 mL cut-off, 95% of samples using a 10 CFU/100 mL cut-off, and 98% of samples using a 100 CFU/100 mL cut-off. ALERT was also assessed for false positives by using concentrated stocks of six non-target bacteria (*Aeromonas, Citrobacter*, *Enterobacter*, *Klebsiella*, *Pseudomonas aeruginosa* and *Serratia*) and for false negatives by using the same non-target bacteria spiked with low levels of *E. coli*. ALERT did not report any false positive values in the absence of *E. coli* and accurately detected *E. coli* in the presence of each of the non-target bacteria, demonstrating excellent specificity and sensitivity. Notably, the ALERT Lab stood out as the only fully automated analyzer, among the 20 tested solutions, that satisfied all the criteria set out in the WHO/UNICEF Target Product Profile (UNICEF, 2019).

## 4 Discussion

### 4.1 ALERT *E. coli* quantification process vs. current standard methods

Conventional culture-based methods for quantifying viable and culturable *E. coli*, such as membrane filtration (MF) with agar plating (ISO, 2014) and various MPN assays (ISO, 1998; ISO, 2012), are based on the assumption of fully homogenized samples and are thus optimized for measuring planktonic bacteria. These approaches are unable to differentiate between free-floating *E. coli* and those aggregated onto fecal or sediment particles. As a result, each particle-associated cluster is counted as a single unit, significantly underestimating the total fecal indicator bacteria (FIB) load (USEPA, 2012b; Jarvis et al., 2010).

In contrast, the ALERT method quantifies all viable and culturable *E. coli* by incubating the entire sample volume with a fluorogenic substrate and monitoring real-time fluorescence during bacterial growth. This whole-sample, growth-based approach—coupled to a quantification mechanism similar in principle to qPCR—yields fluorescence detection times directly proportional to the logarithm of the initial bacterial concentration (Angelescu et al., 2019). Crucially, ALERT overcomes the limitations of MF and MPN by detecting both planktonic and aggregate-bound *E. coli*, as the latter dissociate through cell division and contribute fully to the fluorescence signal (Angelescu et al., 2024a). Planktonic counts can also be independently assessed via sample pre-filtration.

### 4.2 Risk-assessment implications: ALERT *vs.* current standard methods

In waters affected by CSOs, inadequately treated WWTP effluent discharge, stormwater, or agricultural runoff, the discrepancy between MF/MPN counts and the actual number of viable microorganisms can be significant. Discrepancies between comprehensive and planktonic *E. coli* counts of up to two orders of magnitude were previously reported in certain irrigation canals and in insufficiently treated WWTP effluent, while urban waterways, such as the Seine River at the Alexandre III bridge in Paris (2024 Olympic venue for triathlon, paratriathlon and marathon swimming events), revealed comprehensive *E. coli* counts that were more than three times higher, on average, than those measured using traditional laboratory methods (Angelescu *et al*., 2024a). The widespread presence of particle-bound FIB in urban rivers has also been highlighted in earlier studies using enzymatic activity assays, although without quantitative resolution (Garcia-Armisen et al., 2009; Passerat et al., 2011).

In the present study, we report differences larger than one order of magnitude between comprehensive and planktonic *E. coli* counts in the Thames river in the aftermath of a CSO, with planktonic fractions, averaged for the duration of the study, accounting for only 27% of the total *E. coli* load in the Thames River, and for 39% in the Seine River. We also provide convincing supporting evidence of the presence of such fecal aggregates, using epi-fluorescence micropscopy in conjunction with species-specific immunostaining and generic DNA staining to distinguish *E. coli* bacteria in the background of non-specific DNA.

In such waters, standard MPN and MF laboratory methods offer a distorted view of the total FIB load and the associated health risks—akin to estimating the weight of an iceberg by considering only its visible tip. By contrast, the ALERT system provides a much more complete assessment, capturing both the comprehensive FIB count and the planktonic fraction, thereby offering a finer and more accurate picture of microbiological water quality.

### 4.3 ALERT metrology evaluation as on-site laboratory replacement

The side-by-side metrological evaluation of the ALERT and MPN methods was designed to specifically compare the two methods’ ability to quantify planktonic *E. coli*. By removing larger aggregates present in the effluent, the comparison minimized systematic discrepancies caused by the standard laboratory method’s limited sensitivity to aggregate-bound bacteria.

The metrological evaluation demonstrated excellent linearity for both ALERT Lab and ALERT System V2, with coefficients of determination close to unity and no need for slope correction. Nonetheless, small but consistent offsets were observed: a slight overestimation for ALERT Lab and a comparable underestimation for ALERT System V2. These discrepancies are likely due to matrix-specific lag time differences during the initial phase of bacterial growth. Importantly, such offsets can be easily corrected through a single-point calibration prior to long-term deployment in new water matrices. The adjustment procedure is outlined in the following section.

The precision of both ALERT Lab and ALERT System V2 was comparable to that of the standard laboratory, with both instruments—as well as the reference laboratories—displaying slightly better precision at higher *E. coli* concentrations. Notably, the precision obtained in this study significantly surpassed that of earlier metrological characterizations (Angelescu et al., 2019), which used raw (unfiltered) WWTP effluent for spiking. This improvement is primarily attributed to the removal of fecal aggregates (to which the laboratory method is insensitive, leading to systematic undercounting), as well as advancements in ALERT hardware (such as the adoption of sterile disposable measurement cartridges in the ALERT System V2) and refinements in the ALERT detection algorithm.

Additionally, ALERT and MPN measurements from the Berlin time-series study on raw, unfiltered samples enabled estimation of the field precision of both the ALERT Lab instrument and the reference laboratory, by analyzing the distribution of differences between duplicate measurements. The resulting precision of ALERT comprehensive measurements was effectively equivalent to that of the laboratory MPN method, and consistent with previous laboratory precision estimates using raw samples (Angelescu et al., 2019).

Together, these laboratory and field metrology results indicate that, once a matrix-specific offset is corrected via a simple single-point adjustment, ALERT technology achieves accuracy and precision levels that are equivalent to those of standard laboratories when assessing waters with homogeneously distributed FIB. ALERT instrumentation therefore could be used interchangeably with laboratory measurement in recreational water quality monitoring applications. In contrast, in samples containing FIB-rich aggregates, laboratory methods are limited in accurately quantifying the particle-attached bacterial load, whereas ALERT can provide both comprehensive and planktonic *E. coli* counts.

#### 4.3.1 Offset removal due to sample temperature and water matrix effects

The offset of +0.03 Log_10_ units observed during the Berlin time-series study between samples analyzed by the laboratory MPN method and those analyzed by ALERT Lab after cold storage (for a duration equivalent to that of the laboratory samples) suggests that the small overestimation offset (+0.12 Log_10_ units) seen in the on-site ALERT measurements may actually be attributed to bacterial mortality during cold storage, rather than to instrument bias.

An alternative explanation is that for the cold-stored samples, the lower starting temperature led to a longer heat-up ramp to reach the 37°C incubation temperature during the ALERT protocol, causing a slight delay in fluorescence detection and thus reducing the offset. While the present study could not distinguish between these two possible explanations, newer versions of ALERT technology address this issue: the ALERT One and ALERT Lab instruments allow manual input of sample temperature, while the ALERT System V2 can automatically measure it using an external sensor, effectively eliminating small potential measurement offsets arising from temperature differences.

The matrix-specific offsets observed between ALERT and laboratory measurements can be corrected through a single-point offset following a simple procedure:

1. **Collect** approximately 2.5 L of water from the matrix of interest.
2. **Filter** the sample through a 5 µm filter to remove larger aggregates. If the native *E. coli* concentration is estimated to be below 1,000 CFU/100 mL, additionally spike the sample with 5 µm-filtered, non-disinfected WWTP effluent to reach a final concentration between 1,000 and 10,000 CFU/100 mL.
3. **Perform** six independent ALERT Lab measurements and six laboratory MPN measurements on separate aliquots from the prepared water sample, after homogenization through vigorous shaking (seven measurements for ALERT System V2).
4. **Calculate** the geometric means µ_ALERT_ and µ_LAB_
5. **Apply** a correction factor *f* = µ_LAB_/µ_ALERT_ to all future raw ALERT measurements in that matrix (for Log_10_-transformed values, apply a constant offset Δ=Log_10_(µ_LAB_)-Log_10_(µ_ALERT_)).

### 4.4 Short- and long-term effects of CSOs on water quality

This study offers valuable new insights into how water quality responds to different STP events in various environments. The Henley-on-Thames case provided a particularly noteworthy observation: after an isolated CSO event, comprehensive *E. coli* counts gradually declined toward planktonic levels with an unsuspectedly long decay time constant of 27 days, suggesting long-lasting persistence of *E. coli*-rich fecal particles—likely due to sediment trapping and gradual resuspension. To our knowledge, this is the first study to examine CSO impacts from the specific lens of aggregated versus planktonic FIB, revealing previously unrecognized long-term pollution effects.

In contrast, water quality in the Berlin Spree canal showed much faster recovery, with baseline concentrations restored within 24 hours. This can be explained by the local character of the CSO discharges in Berlin and by the limited volume of water in the Canal being rapidly flushed after heavy rainfall events. In the Ablon-sur-Seine case, detailed high-frequency pollutographs recorded a 48-hour return to baseline after storm events. However, given the slow dynamics of aggregate-bound *E. coli* recorded in the Thames, and the increasing frequency of climate-driven storms, it remains uncertain whether such soft-bottom rivers can reliably achieve a true dry-weather baseline between sewage pollution events.

### 4.5 Implications for drinking water treatment and monitoring

This study confirms the widespread and substantial presence of aggregate-bound *E. coli* in major European rivers such as the Seine and the Thames, the results further highlighting the increased resilience of these aggregate-attached bacteria to potabilization treatments—including flocculation, filtration, and chlorine disinfection—which are core treatment steps used in water treatment plants supplying Paris, London, and numerous smaller communities along these rivers. Notably, we observed significantly poorer abatement efficacy—by a factor of 3.9—for aggregate-bound *E. coli* compared to planktonic counterparts. Currently-used MPN and MF culture-based laboratory methods are largely insensitive to aggregate-bound bacterial loads, underscoring the critical importance of measuring comprehensive *E. coli* counts to ensure the safety and reliability of drinking water supplies sourced from such surface waters.

These findings carry even deeper implications for LMICs, where disinfected water supplies and water quality laboratories are often unavailable, and in certain countries most water sources test positive for *E. coli*. In such contexts, point-of-use disinfection solutions—such as the P&G treatment evaluated in this study or equivalent products—can offer a pathway to safe drinking water, but often at relatively high cost, a major limitation in resource-constrained settings. The results presented here demonstrate that portable rapid microbiology analyzers can be effectively used onsite to identify the minimum treatment dose that consistently delivers safe drinking water for a specific contaminated source, even in the presence of fecal aggregates. In the case of the Seine River, for example, our analysis determined that a 50% treatment dose was sufficient, enabling a two-fold reduction in treatment costs. For less-contaminated sources further dose reduction could be possible, significantly improving the cost effectiveness of point-of-use treatment solutions without compromising safety.

When no treatment solution is available, selecting the safest available water—by identifying sources with the lowest *E. coli* concentration—remains the only viable strategy to reduce waterborne disease mortality (UNICEF, 2019). This challenge has led to the development of a semi-quantitative *E. coli* risk level classification system to support risk-based water safety assessments: Compliant: 0/100 mL; Low risk: 1–10/100 mL; Intermediate risk: 10–100/100 mL; High risk: 100–1,000/100 mL; Very high risk: >1,000/100 mL (WHO, 2022b). However, the limitations of laboratory culture-based methods in detecting aggregate-bound bacteria can lead to significant underestimation of total *E. coli* levels, resulting in risk class misclassification and undermining the accuracy of risk assessments by failing to capture the full infectious potential of aggregated FIB and related pathogens. Additionally, the presence of *E. coli* aggregates introduces inherent sample-to-sample variability due to the uneven distribution of particulate matter, further complicating reliable water quality evaluations.

Importantly, the ALERT system’s ability to match the risk class provided by the MPN method in 87% of tests under UNICEF’s Rapid Water Quality Testing initiative speaks to its accuracy and practical utility in LMIC settings. Crucially, in cases where discrepancies occurred, ALERT consistently indicated a higher risk class than the corresponding MPN result—a finding of particular relevance. Based on the evidence presented in this study, it is highly likely that these occasional discrepancies stemmed from the undercounting of particle-bound bacteria by the laboratory rather than overcounting by ALERT. This is because ALERT was configured to measure comprehensive *E. coli* counts (including aggregate-associated bacteria), while the test samples used in validation were spiked with unfiltered wastewater effluent containing both free-floating and aggregate-bound *E. coli*—the latter largely escaping detection by the conventional MPN laboratory method.

### 4.6 Implications for monitoring aquatic sports and recreational waters

Recreational activities in surface waters, widely recognized for their psychological and wellness benefits, have experienced a surge in popularity following the COVID-19 lockdowns (Overbury *et al*., 2023). Efforts to reclaim urban waters for public use and recreation have been further catalyzed by high-profile global events, such as the 2024 Paris Olympics, which hosted aquatic competitions in the Seine River. As a legacy of the Games, the City of Paris plans to open multiple urban river locations for public swimming starting in July 2025, highlighting the cultural and recreational significance of open-water activities.

The ability to deliver accurate, rapid, and frequent testing results is essential for ensuring water safety at recreational bathing and aquatic sports venues. The Berlin time-series case study demonstrates that ALERT measurements achieve a high index of agreement when compared with laboratory data, supporting the use of ALERT as a laboratory substitute under the USEPA site-specific alternative recreational criteria guidelines (USEPA, 2014). Furthermore, when evaluated against the European GM and STV criteria, the observed agreement indicated excellent consistency between the MPN and ALERT methods in regulatory water quality classification. Similar agreement is obtained under the World Triathlon “sufficient quality” STV criterion (<1000 *E. coli* / 100 mL) that is generally applied by for competitive events (including at the 2024 Paris Olympics).

In the Berlin case study, the rapid comprehensive ALERT measurements successfully identified three major contamination events. Under an active management framework, this would have triggered temporary beach closures on those specific days, excluding the corresponding measurements from the long-term water quality classification. As a result, the Berlin Spree Canal could have maintained a “good” status under the European bathing water directive (EU, 2006), rather than being downgraded to “insufficient”—a classification that could ultimately lead to permanent site closure.

In both Paris studies (Ablon-sur-Seine and Alfortville), bathing water quality was classified as “insufficient” under the European bathing water directive (EU, 2006) based on all available methods (ALERT planktonic and comprehensive counts, and approved laboratory MPN counts). While the Alfortville study, conducted during the rainy season without wastewater disinfection, is less representative of summer conditions, the Ablon-sur-Seine study—performed in summer further upstream—reached the same classification. Notably, the Ablon-sur-Seine results highlight the critical value of high-frequency monitoring: while weekly lab and daily ALERT measurements showed similar geometric means, lab data missed the rain-driven STP spikes, underestimating variability (SD: 0.23 Log₁₀) compared to ALERT’s higher variability (SD: 0.40 Log₁₀). This translates into a significantly higher 90th percentile, making it much harder to achieve “sufficient” status under EU criteria.

The Henley-on-Thames case study highlights a striking regulatory discrepancy: while planktonic *E. coli* counts classify the site as having excellent water quality, comprehensive counts—heavily influenced by long-term CSO-driven fecal aggregate pollution—place it in the “insufficient quality” category. Such gaps are critical, as they can create a false sense of safety for water users, leaving them potentially exposed to risks from fecal particles carrying individual FIB loads that cannot be accurately quantified using the current approved laboratory methods.

### 4.7 Enabling citizen science and supporting LMIC water quality monitoring

This study provides strong evidence that ALERT technology offers a scientifically robust, operationally feasible, and economically viable solution for decentralized microbiological water quality monitoring. ALERT One is a recent addition to the ALERT instrumentation line, that was specifically designed to address the operational challenges of remote and resource-limited environments. ALERT One is capable of single-sample analysis without reliance on laboratory infrastructure, stable electricity, or internet connectivity. Its compact, self-contained design, compatibility with solar-powered rechargeable power banks, and reliance on low-cost, long-shelf-life reagents make it well-suited for field deployment in LMICs, where centralized laboratory testing is often inaccessible or unreliable.

The relevance of ALERT One extends well beyond LMIC contexts. Its ability to produce near real-time, quantitative microbiological data—by non-specialists and in the field—has significant implications for citizen science and participatory environmental monitoring. In the Henley-on-Thames case study, citizen scientists independently operated ALERT One devices to generate high-frequency microbiological time series. Such community-generated datasets can provide unprecedented spatial and temporal resolution in microbiological water monitoring—capabilities that are currently lacking even in most high-income countries.

The generation of such high-resolution "microbiological big data" can fundamentally shift our understanding of pollution patterns and their consequences. As demonstrated in the various case studies presented, ALERT technology enables the detection of rapid contamination pulses, evaluation of post-event recovery timelines, and the identification of long-term persistence of fecal contaminants, particularly those associated with aggregated bacteria. These insights are critical for developing improved risk assessment frameworks, enabling predictive models of water quality under extreme weather or infrastructure failure scenarios, and informing the design of adaptive management strategies and regulatory thresholds.

Furthermore, ALERT One facilitates community-level engagement in water safety governance by enabling direct evaluation of water sources and point-of-use treatment efficacy. In this study, ALERT was successfully used to assess the performance of a commercial water purification product (P&G Purifier of Water), demonstrating how field-appropriate microbiological testing could guide cost optimization and informed decision-making at the household or community scale.

From a policy and economic perspective, ALERT One meets UNICEF’s cost target of under $6 per test (in batches of 1,000), far below the cost of commercial laboratory-based E. coli testing in high-income countries, which can exceed $100 per sample with delays of several days. By eliminating logistical bottlenecks, minimizing human error, and enabling automated report generation and data centralization, ALERT One supports high-frequency monitoring necessary to track the dynamic nature of fecal contamination in real-world conditions.

The findings of this study support the broad utility of ALERT technology across both low- and high-resource settings. It enables scalable, rapid-response water quality monitoring, facilitates community involvement and data democratization, and opens the door to a new era of high-frequency microbiological datasets that can inform science, policy, and water management practices.

## 5 Conclusion

This study provides robust validation of the ALERT rapid microbiology technology as a powerful and versatile tool for modern water quality monitoring. Through extensive metrological evaluations, high-frequency field deployments, and real-world case studies across multiple geographic and environmental settings, we demonstrate that ALERT instruments offer accuracy and precision on par with conventional laboratory methods while addressing their critical blind spots and enabling fully-automated rapid field measurements.

Notably, ALERT technology’s ability to quantify both planktonic and aggregate-bound *E. coli* can reveal microbiological risks that standard culture-based methods underestimate due to their intrinsic limitations, with major implications for drinking water safety, recreational water management, and public health risk assessments. By delivering rapid on-site comprehensive measurements, ALERT enables informed, near real-time decision-making, which is especially crucial for actively managing contamination events, optimizing point-of-use treatment strategies, and reducing long-term exposure risks.

ALERT One is a new handheld instrument that was designed specifically for deployment in low-resource settings, where laboratory infrastructure, trained personnel, and reliable supply chains are often lacking. Its successful performance, including in citizen science-led campaigns, underscores its accessibility, operational simplicity, and transformative potential to empower communities, first responders, NGOs, and public health stakeholders with field microbiology capabilities. ALERT technology positions itself as a cost-effective scalable solution not only for LMICs and humanitarian contexts, but also for high-income countries.

The findings presented in this work point to a paradigm shift in microbial water quality monitoring: from slow, lab-bound, and incomplete measurements toward rapid, field-deployable, and scientifically robust solutions. By capturing the true microbial landscape—including often-overlooked aggregate-bound contaminants—ALERT technology offers a critical advance to safeguard public health and build global resilience against waterborne threats.

## 7 Author Contributions

DEA coordinated the manuscript preparation and all scientific activities on behalf of Fluidion, developed the sampling protocols for the Seine River work, and performed the related image processing, data analysis, and interpretation. DEA also coordinated activities with the Henley-on-Thames team. DAS led the instrumentation development, providing technical support and upgrades, and coordinated the related engineering work. EV and MH carried out daily sampling from the Seine River, conducted sample filtrations, and performed the laboratory microbiology work; additionally, EV developed data visualization scripts and performed bacterial staining and microscopy, while MH implemented the P&G disinfection protocols. FK was responsible for sampling, sample preparation, and spiking procedures for the Berlin side-by-side study. NC coordinated the Digital Water City project and the associated activities in Berlin. WS developed the side-by-side sampling protocol for the Berlin studies, conducted data collection and analysis, and drafted the corresponding Materials and Methods sections. All authors reviewed and provided detailed input on the final manuscript.

## 8 Funding

This work was partially supported by the Digital Water City project, with funding from the European Union’s H2020 Research and Innovation Programme under Grant Agreement No. 820954. The development of the ALERT One analyzer was partially supported by NIFA/USDA SBIR Grant

2019-33610-29766.

## 9 Acknowledgments

We acknowledge Omar Bach-Rais, Kai Hong, Patrick Ea, Anna Lemarquand, Mateus Brehmer, Andreas Hausot and Vaizanne Huynh for field work and technical contributions on different parts of the project; Sofia Housni (SIAAP) for facilitating the Ablon-sur-Seine installation and providing the side-by-side laboratory data; David Wallace and Chris Szweda for implementing and coordinating the 2025 Thames River citizen science *E. coli* testing campaign; Philippe Guyot (Board Member in the Association for the Development of Interior Navigation – ADNI), for implementing the 2024 Seine River citizen science *E. coli* testing campaign.

## 10 Supplementary Material

## 11 Data Availability Statement

Data that support the findings of this study are available within the paper or upon request. The 2024 Seine data was provided courtesy of the Fluidion^®^ Open Data Initiative (https://fluidion.com/references/open-data-initiative/2024-seine-water-quality#opendata). Historical rainfall in Paris (measured at the Montsouris station) was obtained from: https://prevision-meteo.ch/climat/mensuel/paris-montsouris. The water quality data collected by the City of Paris and used by the International Olympic Committee during the 2024 Olympics was obtained from: https://www.paris.fr/pages/meteo-de-la-seine-quelle-est-la-qualite-de-l-eau-du-fleuve-27467. Historical rainfall in Henley was obtained from: https://www.visualcrossing.com/weather-history/henley%20on%20thames, whereas reported CSO event duration data for the nearest WWTP (Wargrave) was obtained from: https://www.sewagemap.co.uk/.

## References

Angelescu, D. E. et al. (2019) Autonomous system for rapid field quantification of Escherichia coli in surface waters. J. Appl. Microbiol. 126, 332–343.

Angelescu, D. E., Abi-Saab, D., Ganaye, R., Wanless, D. & Wong, J. (2024a) Addressing underestimation of waterborne disease risks due to fecal indicator bacteria bound in aggregates. J. Appl. Microbiol. 135, lxae280.

BBC (2023) “Dozens fall ill after Sunderland triathlon, health chiefs confirm” https://www.bbc.com/news/uk-england-tyne-66421422. Date accessed: February 2025.

BBC (2024) “Belgium out of triathlon relay as athlete falls ill” https://www.bbc.com/sport/olympics/articles/cgerrl2w19ko. Date accessed: February 2025.

Cabelli, V. J. (1983) Health Effects Criteria for Marine Recreational Waters, EPA 600/1–80-031, U.S. Environmental Protection Agency.

DeFlorio-Barker, S., Wing, C., Jones, R. M. & Dorevitch, S. (2018) Estimate of incidence and cost of recreational waterborne illness on United States surface waters. Environ. Heal. 17:3.

Drechsel, P., Qadir, M. & Galibourg, D. (2022) The WHO Guidelines for Safe Wastewater Use in Agriculture: A Review of Implementation Challenges and Possible Solutions in the Global South. Water 14, 864.

Dufour, A. P. (1984) Health Effects Criteria for Fresh Recreational Waters, EPA-600/1–84-004, U.S. Environmental Protection Agency.

EU (2006) “Directive 2006/7/EC of the European Parliament and of the Council of 15 February 2006 Concerning the Management of Bathing Water Quality and Repealing Directive 76/160/EEC”, Available at: https://eur-lex.europa.eu/eli/dir/2006/7/oj/eng. Date accessed: May 29, 2025.

Fluidion (2024) 2024 Olympics - Seine Water Quality. Available at: https://fluidion.com/references/open-data-initiative/2024-seine-water-quality (Accessed May 10, 2025).

Garcia-Armisen, T. & Servais, P. (2009) Partitioning and fate of particle-associated E. coli in river waters. Water Environ. Res. 81, 21–28.

Greenwood, E. E. et al. (2024) Mapping safe drinking water use in low- and middle-income countries. Science 385, 784–790.

Harder-Lauridsen, N. M., Kuhn, K. G., Erichsen, A. C., Mølbak, K. & Ethelberg, S. (2013) Gastrointestinal Illness among Triathletes Swimming in Non-Polluted versus Polluted Seawater Affected by Heavy Rainfall, Denmark, 2010-2011. PLoS ONE 8, e78371.

IPCC (2021). Intergovernmental Panel on Climate Change - Climate change 2022: Impacts, adaptation, and vulnerability. Working Group II Contribution of to the Sixth Assessment Report of the Intergovernmental Panel on Climate Change.

ISO (1998). International Organization for Standardization—ISO 9308–3 water quality— Enumeration of *Escherichia coli* and coliform bacteria—Part 3: miniaturized method (Most Probable Number) for the detection and enumeration of *E. coli* in surface and waste water, https://www.iso.org/standard/20878.html. Date accessed: February 2025.

ISO (2012). International Organization for Standardization—ISO 9308–2 water quality— Enumeration of *Escherichia coli* and coliform bacteria—Part 2: most probable number method, https://www.iso.org/standard/52246.html. Date accessed: February 2025.

ISO (2014). International Organization for Standardization—ISO 9308–1 water quality— Enumeration of *Escherichia coli* and coliform bacteria—Part 1: membrane filtration method for waters with low bacterial background flora, https://www.iso.org/standard/55832.html. Date accessed: February 2025.

Jarvis, B., Wilrich, C. & Wilrich, P. -T. (2010) Reconsideration of the derivation of Most Probable Numbers, their standard deviations, confidence bounds and rarity values. J. Appl. Microbiol. 109, 1660–1667.

Kay, D. et al. (2004) Derivation of numerical values for the World Health Organization guidelines for recreational waters. Water Res. 38, 1296–1304.

Land, K. J., Boeras, D. I., Chen, X.-S., Ramsay, A. R. & Peeling, R. W (2019) REASSURED diagnostics to inform disease control strategies, strengthen health systems and improve patient outcomes. Nat. Microbiol. 4, 46–54.

Marshall, K. E. et al. (2020) Lessons Learned from a Decade of Investigations of Shiga Toxin– Producing Escherichia coli Outbreaks Linked to Leafy Greens, United States and Canada. Emerg. Infect. Dis. 26, 2319–2328.

Overbury, K., Conroy, B. W. & Marks, E. (2023) Swimming in nature: A scoping review of the mental health and wellbeing benefits of open water swimming. J. Environ. Psychol. 90, 102073.

Passerat, J., Ouattara, N. K., Mouchel, J.-M., Rocher, V. & Servais, P. (2011) Impact of an intense combined sewer overflow event on the microbiological water quality of the Seine River. Water Res. 45, 893–903.

Ramírez, S. B., Meerveld, I. van & Seibert, J. (2023) Citizen science approaches for water quality measurements. Sci. Total Environ. 897, 165436.

Sewak, P., 2020. An Impact Assessment of Safe Water Interventions in 154 Communities in Telangana State, Safe Water Network. Available at: https://safewaternetwork.docsend.com/view/nym6wm3r24vnshaz. Date accessed: February 2025.

SimpleLab, 2025. Coliform Enumeration Surface Water Test, Available at : https://gosimplelab.com/shop/d7987a56-4179-4c4f-9f9c-ea34fa8403cd. Date accessed: February 2025.

Spalding, A., Goodhue, R. E., Kiesel, K. & Sexton, R. J. (2023) Economic impacts of food safety incidents in a modern supply chain: E. coli in the romaine lettuce industry. Am. J. Agric. Econ. 105, 597–623.

UNICEF (2019) UNICEF Target Product Profile – Rapid *E. coli* Detection Tests, Version 3.0, Available at: https://www.unicef.org/supply/media/2511/file/Rapid-coli-detection-TPP-2019.pdf Date accessed: May 10, 2025.

UNICEF (2023) Triple Threat: How disease, climate risks, and unsafe water, sanitation and hygiene create a deadly combination for children. ISBN: 978–92-806-5438-7.

UNICEF (2025) Rapid Water Quality Testing, Available at: https://www.unicef.org/innovation/rapid-water-quality-testing Date accessed: June 5, 2025

UNICEF-WHO (2023) Progress on household drinking water, sanitation and hygiene 2000-2022: Special focus on gender. ISBN 978–92-806-5476-9.

USEPA (2009). U.S. Environmental Protection Agency—Method 1603: *Escherichia coli* (E. coli) in water by membrane filtration using modified membrane-thermotolerant *Escherichia coli* agar (Modified mTEC). https://19january2017snapshot.epa.gov/sites/production/files/2015-08/documents/method_1603_2009.pdf. Date Accessed: February 2025.

USEPA (2012b) U.S. Environmental Protection Agency - Recreational Water Quality Criteria, USEPA Report 820-F-12-058.

USEPA (2014) U.S. Environmental Protection Agency - Site-specific alternative recreational criteria technical support materials for alternative indicators and methods. EPA-820-R-14-011.

USEPA (2021) U.S. Environmental Protection Agency - Alternative Methods Calculator Tool User Guide. EPA 821-B-21-001.

Walt, V. (2023) Inside the Billion-Dollar Effort to Clean Up the World’s Most Romantic River. Time Magazine, https://time.com/6261729/seine-clean-up-paris-olympics-2024/ Date accessed February 2025.

WHO (2021) World Health Organization - Progress on household drinking water, sanitation and hygiene 2000-2020: five years into the SDGs. ISBN 978–92-4-003084-8.

WHO (2022a) World Health Organization - State of the World’s Drinking Water: an urgent call to action to accelerate progress on ensuring safe drinking water for all. ISBN 978–92-4-006080-7.

WHO (2022b). World Health Organization - Guidelines for Drinking Water Quality, Fourth Edition incorporating the first and second addenda. ISBN 978–92-4-004506-4.

WHO-UNICEF (2022) World Health Organization and UNICEF Joint Monitoring Programme for Water Supply, Sanitation and Hygiene (JMP) - Laboratory assessment: Fluidion ALERT Lab. https://washdata.org/reports/laboratory-assessment-fluidion-alert-lab Date Accessed: February 2025.

WHO (2025) World Health Organization – Water Sanitation and Health: International Scheme to Evaluate Household Water Treatment Technologies – Products Evaluated. https://www.who.int/tools/international-scheme-to-evaluate-household-water-treatment-technologies/products-evaluated. Date Accessed: February 2025.

